# Assessing the accuracy of paired and random sampling for quantifying plant–plant interactions in natural communities

**DOI:** 10.1101/2022.08.09.503341

**Authors:** Richard Michalet, Gianalberto Losapio, Zaal Kikvidze, Rob W. Brooker, Bradley J. Butterfield, Ragan M. Callaway, Lohengrin A. Cavieres, Christopher J. Lortie, Francisco I. Pugnaire, Christian Schöb

## Abstract

Interactions among plant species in extreme ecological systems are often inferred from spatial associations and quantified by means of paired sampling. Yet, this method might be confounded by habitat-sharing effects, in particular when microenvironmental heterogeneity and stress are high. Here, we address whether paired and random sampling methods provide similar results at varying levels of environmental heterogeneity. Furthermore, we investigate how the relationship between species preferences and abiotic severity influences the outcome of these two methods. We quantified spatial associations with the two methods at three sites that encompass different micro-environmental heterogeneity and stress levels: semi-arid environments in Canary Islands, Spain and Sardinia, Italy and a cold alpine environment in Hokkaido (Japan). Then, we simulated plant communities with different levels of species micro-habitat preferences, environmental heterogeneity and stress levels. We found that differences in species associations between paired and random sampling were indistinguishable from zero in our model simulations. At each site, there were strong differences between beneficiary species in their spatial association with benefactor species, and associations became more positive with increasing stress in Spain. Most importantly, there were no differences in the results yielded by the two methods at any of the different stress levels at the Spanish and Japanese sites. At the Italian site, although micro-environmental heterogeneity was low, we found weakly significant differences between methods that were unlikely due to habitat-sharing effects. We conclude that the paired sampling method can provide significant insights into net, long-term effects of plant interactions in spatially conspicuous environments.

## INTRODUCTION

The interplay between competition and facilitation in shaping plant communities has been intensively studied for more than three decades (Goldberg and Barton 1992; Callaway et al. 2002; Losapio et al. 2021). Most empirical studies assessing these two ecological processes in natural communities have focused on the net outcome of pairwise interactions by comparing the performance of a target species (or dependent community) with and without neighbours often by means of pairwise comparison. This method, sometimes called the observational method (Kershaw and Looney 1964; Maestre et al. 2005), compares the performance of the target species between spatially proximate locations within the same community that are naturally with and without neighbours. It avoids artificial modifications of environmental conditions that could confound the results. However, it can only be applied in situations where the clear and conspicuous presence and absence of a target species (i.e., nurse or competitor) can be determined unambiguously, and this method would be preferable in habitats where vegetation cover is patchy (Liancourt and Dolezal 2020). Consequently, this method has been used mostly for assessing facilitation rather than competition because facilitation has been described more frequently in conditions of low biomass and vegetation cover (Cavieres et al. 2014).

Two sampling methodologies have been used for the observational method. The random sampling method is based on unique set of randomly chosen locations sampled within the same area, some of which contain plants and some not (Kikvidze et al. 2005). On the other hand, the paired sampling method involves deliberately identifying pairs of samples, one randomly including a nurse plant (or a competitor) or vegetation patch and a random, nearby sample from an adjacent area where the facilitator or competitor species under study is absent (Cavieres et al. 2014). The facilitator and beneficiary species can be associated in space in the absence of interactions (i.e., the “oasis effect” often observed in arctic or alpine ecosystems, Muc et al. 1989) if both share the same abiotic requirements, such as deeper soil patches or rock shelters. Consequently, important requirements of the method are that sampled vegetation patches and open areas must be located in similar microhabitats. However, the paired sampling method may be statistically biased towards over-estimating facilitation (Steinbauer et al., 2016). A point of argument is that microhabitat heterogeneity may lead to facilitation overestimation. Nevertheless, it is important to keep in mind that environmental heterogeneity is itself created by long-term feedback mechanisms between vegetation and the abiotic environment (Bera et al. 2021).

Ecosystem-engineering effects generate positive, reciprocal feedback processes “that operate by modifying any of several features of the environment, including water, pH, soil elements, light, temperature, wind, fire, or allelopathic toxins” (Wilson and Agnew 1992). These biophysical interactions then produce a stable vegetation mosaic, with environmental conditions now differing between areas with and without vegetation in what otherwise was previously a uniform environment. Yet, it is important to note that, if microenvironmental heterogeneity within plant communities is biotically driven through long-term ecosystem engineering, they should not be considered as confounding effects but as an outcome of plant interactions.

Consequently, there is a need for the paired sampling method to focus on accounting – if possible – for what might be termed pre-existing (i.e., pre-vegetation development) small scale environmental heterogeneity and eventually disentangling that component from ecosystem-engineering effects. Addressing this point will enable us to draw important conclusions from the results of paired sampling studies. For example, Michalet et al. (2015a) and Noumi et al. (2016) proposed disentangling short-from long-term effects of neighbours using both the paired sampling and removal procedures in the same community. They argued that short-term effects could be quantified using the removal method (with neighbours vs. removed-neighbours conditions) and long-term effects by comparing target responses in removed-neighbours vs. naturally open conditions without neighbours (Noumi et al. 2016). Thus, the net neighbour effects (i.e., the sum of short- and long-term effects) are those quantified by the paired sampling method.

The crucial point is how to separate pre-existing microenvironmental heterogeneity from ecosystem-engineering effects. This is important because both pre-existing spatial environmental heterogeneity and facilitation (stress-gradient hypothesis, Bertness and Callaway 1994, hereafter SGH) are expected to increase with increasing environmental harshness (Steinbauer et al. 2016). Co-analyzing environmental factors, as suggested by Steinbauer et al. (2016), is one route to address this problem, but it is not always easy to determine whether differences between vegetated and bare patches are pre-existing or due to ecosystem engineering effects, as explained above. However, we suggest that by focusing on conspicuous micro-topographic variations that occur in both vegetated and bare patches within the same community, ecologists can account for the likely pre-existing environmental heterogeneity using the paired sampling method, thus overcoming its drawbacks.

In order to explore this tenet, we first used a modeling approach assessing the effects of different environmental and biotic factors on spatial association as quantified with the paired and random sampling methods. In particular, we tested the role of within-community habitat heterogeneity, abundance and ecological preferences of nurse (i.e., the potential facilitator species) and beneficiary species (i.e., the species facilitated by the nurse). We also applied these two methods (i.e., random and paired sampling methods) in three different real-world ecosystems subjected to varying stress levels and exhibiting contrasting soil heterogeneities. We posed two main questions in both the modeling study and the field systems: i) are spatial associations detected with the two methods significantly different when preexisting within-community environmental heterogeneity is high? ii) are spatial associations detected with the two methods affected by species preferences and stress level?

## MATERIALS AND METHODS

### Modeling study

The plant community simulation consisted of two modeling approaches. The first modelling approach assumed a homogeneous space, while the second modelling approach used a gradient over a spatial grid. Our rationale, here, is to simplify the community including few, basic biotic components and to focus on the differences between the two methods. We do not aim at reproducing every and each community assembly process or spatial pattern.

In the homogeneous space model (Wiegand and Moloney 2014), we simulated a plant community composed of nurse (α) and beneficiary (β) plant species. We considered the following abiotic and biotic factors: (i) habitat quality (q), (ii) nurse abundance (i.e., landscape cover) (a_α_), (iii) beneficiary abundance (i.e., landscape cover) (a_β_), (iv) nurse habitat preference (h_β_), (v) beneficiary habitat preference (h_β_), (vi) beneficiary affinity to nurse (β α). A given plant community was characterized by a set of spatial units x (n = 500^2^). We adopted a presence–absence assembly model where the occurrence y of nurse plants α and beneficiary plants β is given by y = 1 for species presence and y = 0 for species absence.

Nurse occurrence is given by 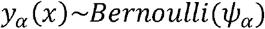where the occurrence probability ψ, of a nurse plant species α given the environmental conditions in x was calculated as the geometric mean of nurse abundance (aα) and ‘habitat suitability’ (hsα), suchthat 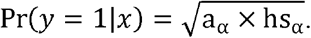The factor ‘habitat suitability’ (hs _α_) was calculated as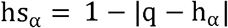, where we considered the differences between habitat quality (q) and nurse habitat preference (h _α_). For instance, a stressful environment would be less unsuitable to a stress-resistant plant rather than to a demanding plant, and vice versa.

Beneficiary occurrence is given by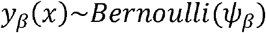, where the occurrence probabilityψ, of a beneficiary plant species β given the conditions in x was calculated as the geometric mean between beneficiary abundance (a_β_) and habit suitability (hs_β_) as well as with a matching factor (m_β_), such that 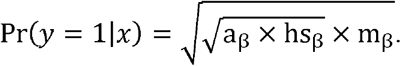Here, the nurse-beneficiary matching factor (m_β_) was calculated as 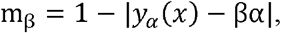where we considered the combination of nurse occurrence (y_α_) and beneficiary affinity to nurse (βα).

Simulation model parameters were set as following: q = 0.0, 0.1, 0.9; a_α_ = 0.050, 0.133, 0.217, 0.300; a_β_ = 0.010, 0.073, 0.137, 0.200; h_α_ = 0.200, 0.467, 0.733, 1.000; h_β_ = 0.200, 0.467, 0.733, 1.000; β α= 0.000, 0.333, 0.667, 1.000. Sampling efforts were equal to 5% and 10% of the community. Each factor combination was replicated in a fully-factorial experiment, resulting in n = 4,096 replicates (communities).

We proceeded with sampling the community according to the random or pairwise approaches. In the random approach, a set of plots is sampled randomly from a uniform distribution, i.e., with constant probability for each spatial unit x. In the pairwise approach, a set of plots is sampled randomly from a uniform distribution of plots in which the nurse is present (i.e., for those x where y_α =_1), while a second set of plots is sampled randomly from a uniform distribution of plots in which the nurse is absent (i.e., for those x where y _α=_0). Pairwise plots are generated from randomly sampling plots where the nurse is present or absent, as opposed to random plot where nurse presence is not considered a priori. Then, we recorded whether the nurse α and the beneficiary β plants were present or absent in each plot for each of the two sampling methods (see also the R code in SI).

We then addressed the influence of sampling methods on the estimated facilitation effects. Facilitation effects were calculated as the dependency of beneficiary species on nurse species using a generalized linear model of the form 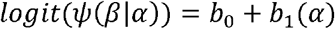 where *b*_0_ (intercept) and *b*_*1*_ (slope) are the parameters estimating the occurrence of beneficiary plants in the absence and presence of nurse plants, respectively. Then, we looked at the impact of abiotic and biotic factors on normalized *b*_*1*_ (i.e., *z-*score) parameters for each sampling method in conjunction (see SI). This was accomplished by running linear models with *z-*score parameters as response and habitat quality, nurse abundance, beneficiary abundance, nurse habitat preference, beneficiary habitat preference, beneficiary affinity to nurse, and sampling effort as predictors, as well as with sampling method as predictor alone and in interactions with all these previous factors (see SI). Finally, we considered the relative differences in facilitation effects *b*_*1*_ between pairwise and random sampling as 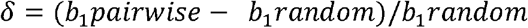. These relative differences δ were then tested for differences from zero using a *t* test, and for the effects of abiotic and biotic factors using a linear model.

In the second approach, we assumed a heterogeneous space simulating communities with a different spatial structure of environmental variation (Buckley et al., 2016). We generated a bivariate spatial pattern with an abundance gradient using the codispersion model proposed by Buckley et al. (2016). We simulated species co-occurrence patterns as a raster of 10,000 5 × 5 grid points over an area of 500 × 500. Species abundance decreased along the *y-*axis for both species. We used this spatial pattern as it previously showed the best codispersion accuracy (Buckley et al. 2016). The simulation created a set of quadrat abundance values *n* for nurse plant and beneficiary plant species. Abundance was generated as 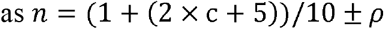sampled from a normal distribution, where *c* is the *y-*axis coordinate and ρ is a random noise introduced to break the ties. Random noise ρ is approximately a fifth and a tenth of mean abundance for the nurse and the beneficiary, respectively (Buckley et al. 2016; see also SI). The simulation pattern was replicated 50 times for each species (see also the R code in SI).

Species in each pattern were sampled either randomly or pairwise. Notice that this second approach differs from the first one in that species, rather than plots, were sampled randomly or pairwise (i.e., the same grid point). The reason is that the codispersion pattern fills the whole spatial grid and has abundance values in each grid point (i.e., the nurse and the beneficiary species are present everywhere with different abundance, that is there are no grid points without nurse plants). Each nurse and beneficiary species pattern was sampled from a uniform distribution 500 times.

For each pattern, we computed the facilitation effect looking at the dependency of beneficiary species on nurse species by means of Generalized Least Square (GLS) models (Pinheiro and Bates 2000). Each GLS fits beneficiary abundance data as response variable and nurse abundance data as predictor. Linear spatial correlation along the *y-*axis was included as error correlation structure. Then, we compared the GLS model parameters (beta and t-value) between the two methods using linear mixed models (Pinheiro and Bates 2000). Two separate models were run for beta and t-value, including spatial pattern replicate as a random effect. Finally, we calculated the type-II (i.e., false negative) error rate for each method by looking at the amount of misidentified, unsignificant (i.e., t-value lower than 1.96) cases.

Simulations and analyses were done in R environment version 4.0.2 (R Core Team 2020).

## Field communities

### Study sites, communities, and species

We aimed to compare the association between nurse and beneficiary species using paired sampling and random sampling methods in different natural conditions of pre-existing environmental heterogeneity and stress. We selected two summer-dry Mediterranean-type systems and an alpine-type system, as dry and cold ecosystems are considered as the most likely to exhibit facilitation effects by nurse plants. The dry site with greater within-community pre-existing environmental heterogeneity due to high physical disturbance was located in the Gran Canaria Island (Spain; Agaete, 28°05’20’’N, 15°42’18”W). A dry site in Sardinia Island (Italy; Dorgali, 40°18’58’’N, 09°32’09”E) and a wet alpine site in Japan were our homogeneous sites (43°40′2″N, 142°55′16″E).

The Spanish site was located at sea level in a climate with a long summer drought (10 months) due to its very low latitude for a Mediterranean climate. Mean annual temperature is 20°C and annual rainfall is 195 mm. The Italian site was located at 180 m a.s.l. with a shorter summer drought (six months) due its higher latitude. Mean annual temperature is 16°C and annual rainfall is 480 mm. The Japanese site was located at 2158 m a.s.l. near Mt. Koizumi (43°40′2″N, 142°55′16″E). Climate is temperate oceanic alpine with mean annual temperature and rainfall of -2°C and 4000 mm, respectively.

At the Spanish site vegetation is an open shrubland dominated by *Euphorbia balsamifera*, a species abundant in the Macaronesian region (South West of Morocco and Canary Islands) (Fig. 1). At our site, the plant cover ranges between 30 and 70% depending on local factors. *Euphorbia* was used both as nurse and beneficiary species, and the other beneficiary species was the forb *Astydamia latifolia*, a perennial species from the Apiaceae family. In general, in the three systems nurse species were chosen because of their dominance and target species because of obvious patterns of association with nurses. At the Spanish site, *Astydamia* cover varied between 5 and 75% depending on local factors modifying stress levels, such as distance to the ocean and exposure.

**Fig. 1.**
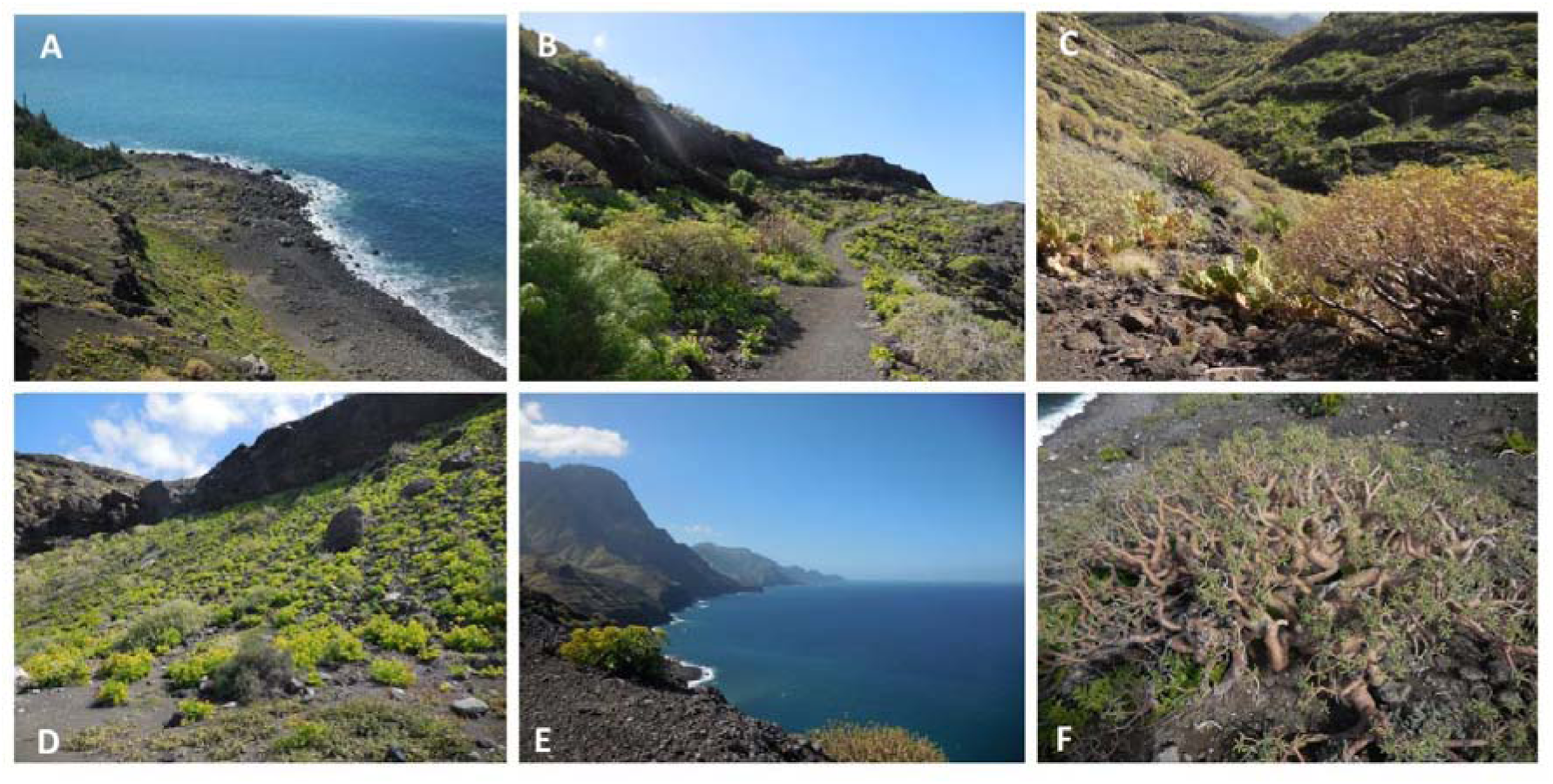
The Spanish site (Gran Canaria): A: the least stressed subsite below the black cliff, B: the intermediately stressed subsite, C: the most stressed subsite on the left part of the slide (in southern exposure, with the intermediately stressed subsite on the right side of the slide in north exposure), D: abundance of *Astydamia* in the open in the least stressed subsite, E: general view of the Atlantic coast at the site with *Astydamia* (left) and *Euphorbia* (right) in open conditions, F: *Astydamia* (in green on the bottom left) below the canopy of *Euphorbia* in the most stressed subsite. Note that in all slides *Astydamia* is very easy to locate thanks to its very light green colour.

At the Italian site, the vegetation is an old *Olea europaea* orchard plantation that was abandoned and now transformed in a savannah community used for grazing by sheep (Fig. 2). Vegetation cover is approximately 50% for the trees, which are regularly spaced due to management. The associated herbaceous community cover is between 20 and 90% depending on microhabitats. It is dominated by grasses and two forb species that were selected as beneficiaries, *Smyrnium rotundifolium* and *Cynara cardunculus*, while the nurse species was *Olea europaea*.

**Fig. 2.**
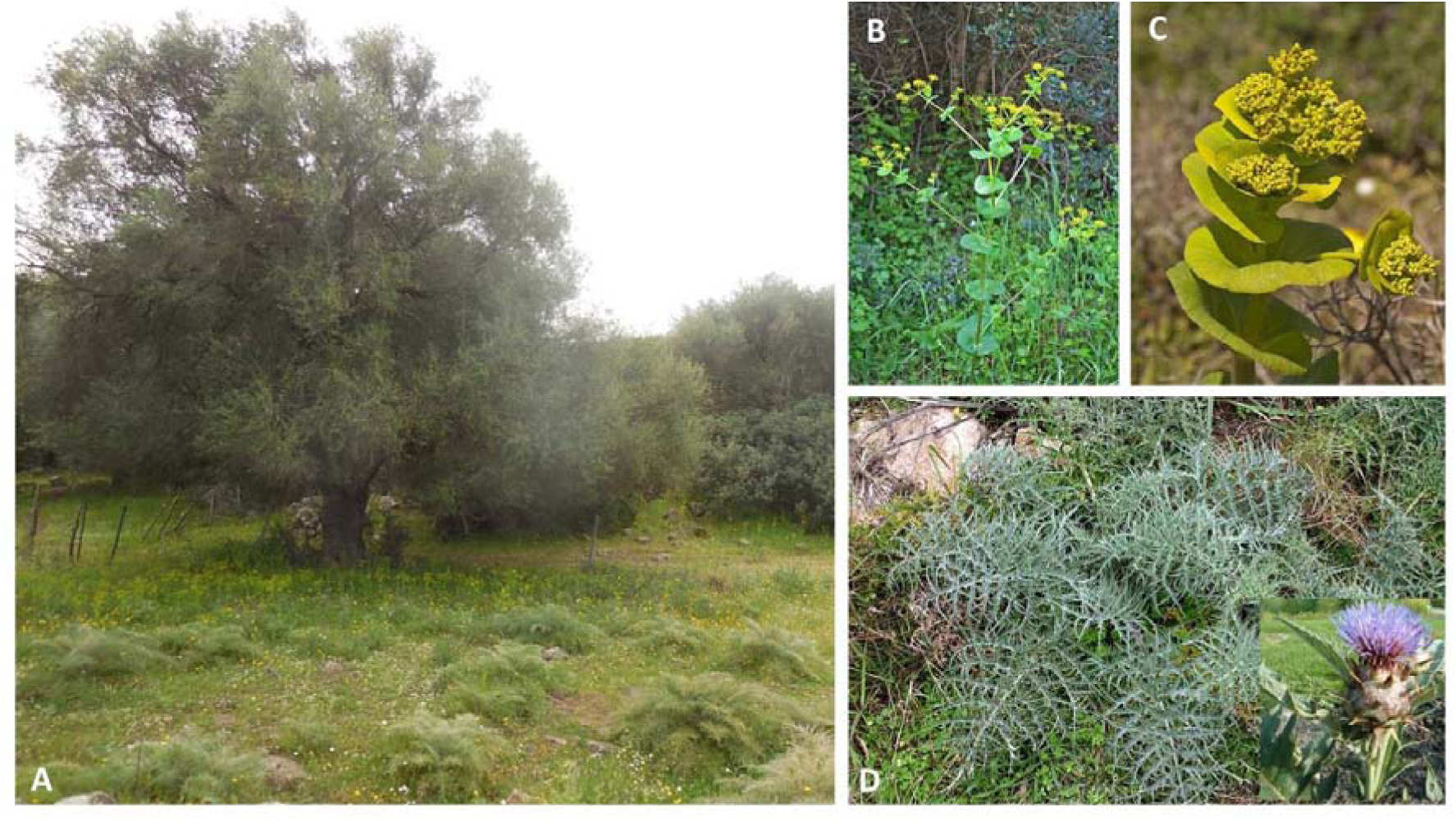
The Italian site (Sardinia): A: the northern position of the canopy of *Olea* with below in green *Smyrnium* and in the open (in the front) *Cynara*, B and C: *Smyrnium*, D: *Cynara*.

The Japanese site was located on a mild slope (5°) near the summit of Mt. Koizumi. The vegetation was patchy, its cover reaching 75%, and was dominated by cushions of *Diapensia lapponica* and lichens. Other prominent species were *Arcterica nana, Carex stenantha var. taisetsunensis, Empetrum nigrum, Loiseleuria procumbens* and *Salix nummularia*. Mean diameter of the cushions of the nurse species *Diapensia* was 35.5 ± 20.3 cm (n = 70). *Diapensia lapponica* was the nurse and the beneficiaries were species richness and *Salix nummularia*.

### Experimental design

In each site we sampled with the two methods -paired and random sampling. At the Spanish site we sampled three very similar communities but having contrasting levels of stress, whereas at both the Italian and Japanese sites we sampled only one community. At the Spanish site we selected three subsites along a complex gradient of increasing stress due to distance to the ocean and exposure. The least stressed subsite had a north-west exposure and was quite close to the ocean (less than 100 m). Thus, the lower drought stress was very likely due to the amelioration of drought by water spray, as found by Forey et al. (2008) in the French coastal dunes (Fig. 1 A). Close to the ocean the cover of *Astydamia* was highest (approximately 75%), with lots of individuals occurring in the open between *Euphorbia*, the latter having the lowest cover (approximately 30%) at this subsite. The intermediately stressed subsite had the same exposure but was farther from the ocean (several hundred meters). The cover of *Astydamia* and *Euphorbia* were both high (approximately 50% for each species), with no obvious spatial patterns of association or repulsion. The most stressed subsite was located at a similar distance from the ocean, but had a south west exposure. The cover of *Astydamia* and *Euphorbia* were the lowest (5%) and highest (70%), respectively, at this site, and most *Astydamia* were only observed below the canopy of the nurse. Physical disturbance was high in all three subsites, with deep ravines alternating with vegetated patches, due to the occurrence of the three subsites in slopes on volcanic scoria.

At the Italian site, pre-existing environmental heterogeneity was very low, since the site was in a dry floodplain on calcareous rock. Topography was flat and soils deep with a fine texture. We chose to assess the association of two understory (beneficiary) species with *Olea* at three canopy positions because there were obvious differences in abundance and cover of the dominant forb *Smyrnium*, between the three *Olea* canopy positions, likely due to higher stress with increasing irradiance. The other beneficiary species, *Cynara*, did not show obvious differences in abundance related to *Olea* canopy position but mostly occurred in the open where its cover can be high (approximately 70%). In contrast, *Smyrnium* was more abundant below trees, and in particular in the northern side of the canopy (approximately 70% cover vs. 5% in the southern side).

At the Japanese site, sampling was performed on alpine scree soils of a volcanic origin in a mature alpine community with a homogeneous matrix of cushions. Thus, the pre-existing environmental heterogeneity was apparently low, with regular action of snow on a nearly flat terrain (Fig. 3).

**Fig. 3.**
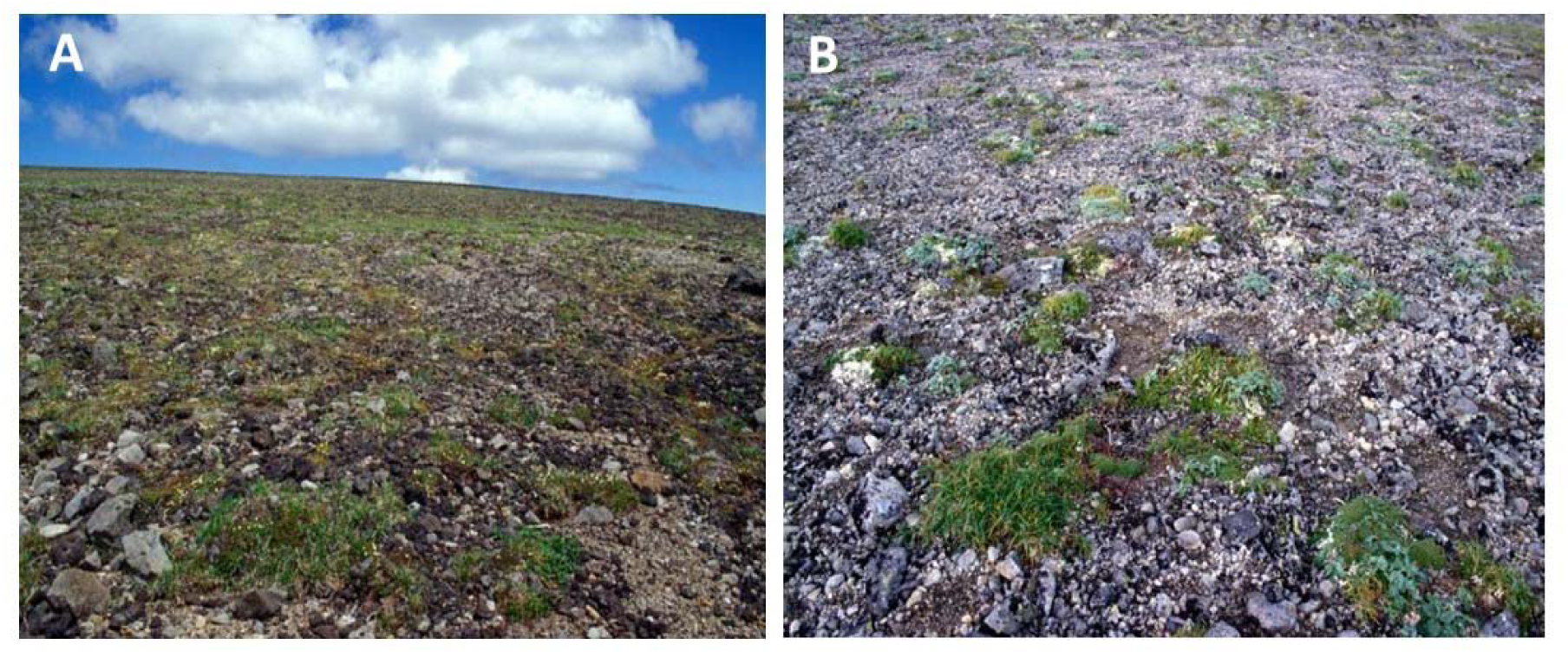
The Japanese site (Hokkaido): A: the sampling site with low apparent pre-existing heterogeneity; B: a typical cushion of *Diapensia lapponica*.

### Vegetation sampling

In the Spanish site, at the three stress levels, and in the Italian site at the three canopy positions, we sampled the performances of beneficiary species with and without neighbours, using both sampling methods. At each of the three subsites in the Spanish site, for the paired sampling method we randomly selected 30 *Euphorbia* patches with a size above 1 m^2^ and positioned a quadrat of 250 cm^2^ on the northern side of the canopy at mid-distance between the trunk and the edge of the canopy. Another quadrat of the same size was randomly positioned in open conditions in vicinity (one meter) to the *Euphorbia* patch and in similar environmental conditions (same microtopography and soil). In the 60 quadrats of each subsite we visually estimated the cover of the two beneficiary species, *Astydamia* and *Euphorbia* seedlings. For the random sampling method, we randomly chose the position of 60 plots and sampled with the same quadrat the cover of the two beneficiary species. The position of each quadrat was determined by throwing the quadrat to the back and walking 10 meters along transects between two quadrats to fully explore the whole community (but see Dudley 1982 for a straightforward random approach). Thus, the number of plots per microhabitat condition at each subsite was dependent on the cover of each microhabitat. It varied between 14 for open conditions to 46 in adult *Euphorbia* patches of the most stressed site.

At the Italian site, for the paired sampling method we randomly selected 30 *Olea* individuals that were spaced at least 30 meters apart from each other. The height and diameter of the canopy of selected trees varied between 3 and 8 m and 4 and 10 m, respectively. For each of the 30 tree individuals and at each position of the tree canopy (north, south and west) we positioned a quadrat of 1 m^2^ at mid-position between the trunk and the edge of the canopy. Then, we positioned the paired quadrats for the open positions at 5 m from the edge of the canopy for each individual. In each quadrat we recorded the number of individuals of the two beneficiary species. For the random sampling method, because the cover of the tree population was approximately 50% and because the size of the trees prevented random throwing of a quadrat, we chose to separately sample tree and open plots. Tree plots and their positions were selected by the same technique as used for the paired sampling approach. For open plots, we first randomly selected 30 open areas between trees with the same technique as the one used to select tree individuals. Then, we randomly threw the quadrat three times in the 30 selected open areas for positioning the three open plots that were systematically affected by treatments (north, west and south). Thus, for the Italian site the number of replicates was the same in all treatments, although plots were different for the two samplings.

At the Japanese site, for the paired sampling method we haphazardly chose 70 cushions of *Diapensia lapponica*, and all plants growing within these selected cushions were identified to the species level and their abundance recorded. We measured the maximum and minimum axes of each cushion to estimate its area. To obtain comparable samples for assessing species richness in surrounding ‘open’ areas (areas not covered by cushions), areas matching the size of each sampled cushion were surveyed at haphazardly selected paired points away from each sampled cushion. Random sampling was performed with a small wire square (10 cm on a side), which was randomly placed 100 times within the sampling area; then all established plants within the square were identified and recorded.

### Statistical analyses

For the Spanish site, the effects of the stress level, method (paired *vs* random), neighbour (*Euphorbia vs* open), beneficiary (*Euphorbia vs Astydamia*) and all their interactions on cover of the target species was tested using a linear mixed effects model. As random terms, we included the replicate for which the two beneficiary species were quantified, and the interactions of replicate with the stress level, stress level and method, and stress level and method and neighbour to account for pseudoreplication.

For the Italian site, the effects of method (paired *vs* random), neighbour (*Olea vs* open), canopy position (three cardinal directions), beneficiary (*Cynara vs Smyrnium*) and all their interactions on abundance of the beneficiary species was tested using a linear mixed effects model. As random terms we included the replicate for which the two beneficiary species were quantified, and the interactions of replicate with method, neighbour and method, canopy position and neighbour and method.

For the Japanese site, the effect of method (paired *vs* random), neighbour (*Diapensia vs* open) and their interaction on species richness was tested using a linear model with square-root-transformed richness data to meet model assumptions. The effect of method, neighbour and their interaction on the frequency of occurrence of the beneficiary species *Salix nummularia* was tested using a generalized linear model with presence/absence of Salix as the response variable and the corresponding binomial distribution of error terms.

The linear mixed effects model analyses were conducted with asreml for R environment version 4.1 (Butler 2020) and the convenience functions for fitting negative variance components and for type-I analysis of variance provided by pascal (Niklaus 2019). Sequential (type-I) testing of factors was justified with our hierarchical experimental designs where the two different methods were assessed with/without neighbour species and different beneficiary species growing with/without neighbour. Therefore, the general sequential order of factors following the pattern “method > neighbour > beneficiary” was followed during testing. Linear models and generalized linear models and the corresponding type-I analyses of variance were conducted with the base functions in R version 4.0.4 (R Core Team 2020).

## RESULTS

### Modeling results

The results of our first-approach simulation indicate no significant differences between random and pairwise methods in estimating facilitation effects (absence of correlation as t = -1.59, P = 0.113). Looking at the effects of abiotic and biotic factors and sampling method on facilitation effect (Table S1), we found that the variance in standardized parameters (z-score) was: (i) marginally explained by nurse habitat preference h_α_ (P=0.061); (ii) significantly explained by beneficiary habitat preference h_β_ (P=0.046); (iii) significantly explained by beneficiary affinity to nurse βα alone and depending on sampling method (P<0.001 and P=0.020, respectively). When considering parameter estimates (Table S2), beneficiary affinity to nurse β α significantly increases facilitation effects quantification, both per se (estimate = 8.02 ± 0.42 SE, *P*<0.001), as reasonably expected, as well as with higher impact using the pairwise method (estimate = 1.38 ± 0.59 SE, *P* = 0.020).

The relative net differences δ in facilitation effects between pairwise and random sampling were neither positive nor negative overall (mean = -0.058, 95%CI = -0.130 – 0.014, Fig. 4A). Among the abiotic and biotic factors we tested, only beneficiary affinity to nurse significantly explained δ(F_1,4088_ = 207, *P* < 0.001). In particular, the higher the beneficiary affinity to the nurse the larger the differences in facilitation effects between the two methods (estimate = 1.38 ± 0.10 SE, *P* < 0.001, Fig. 4G). In synthesis, although our model shows that there were no significant differences between the two methods overall, the paired sampling as compared to the random one tended to provide higher parameter estimates of facilitation effect with increasing beneficiary preference for the nurse.

**Fig. 4.**
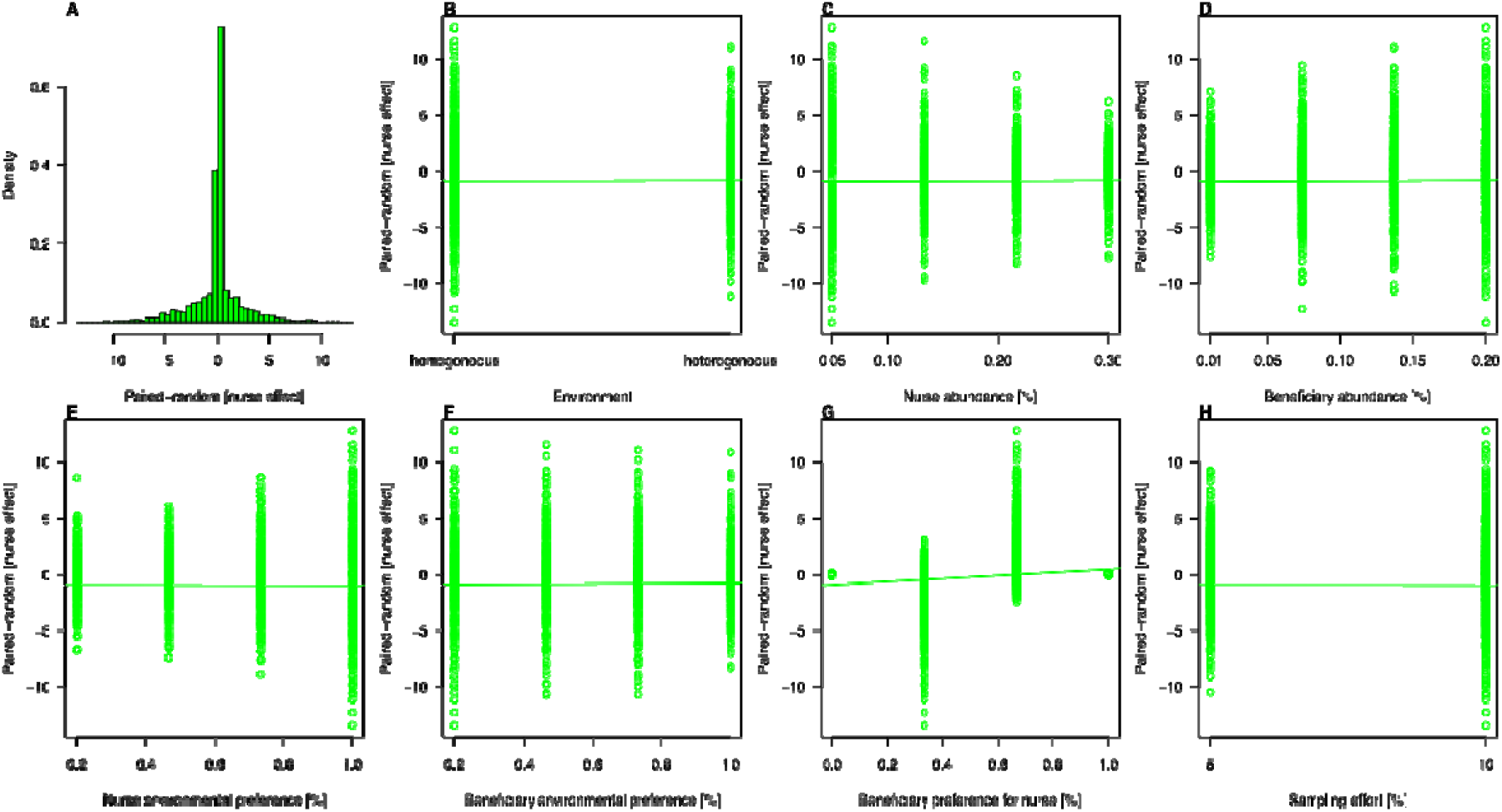
(A) Distribution of relative differences in facilitation effects between pairwise and random sampling. (B–H) Impact of abiotic and biotic factors on Lines represent the linear regression model.

The second-approach simulation, that of species codispersion over a decreasing abundance gradient, yielded qualitatively similar results. Results indicate that both sampling methods are valid for identifying facilitation. Yet, the two methods differ quantitatively as the pairwise method produced significantly higher betas (11.3%, *P*=0.021) that were closer to one and higher t-values (65.1%, *P*<0.001) than random ones (Fig. 5). Furthermore, the variance of dependency parameter estimates was much higher for the random sampling (estimate sd = 0.30, t-value sd =108.4) than for the pairwise sampling (estimate sd = <0.01, t-value sd = 12.6). Finally, while random sampling sometimes failed at correctly detecting facilitation (four false negative cases, type-II error rate of 8%), the pairwise method was correct over all 50 simulations. In summary, the two methods are qualitatively similar, but the pairwise method is more precise and accurate.

**Fig. 5.**
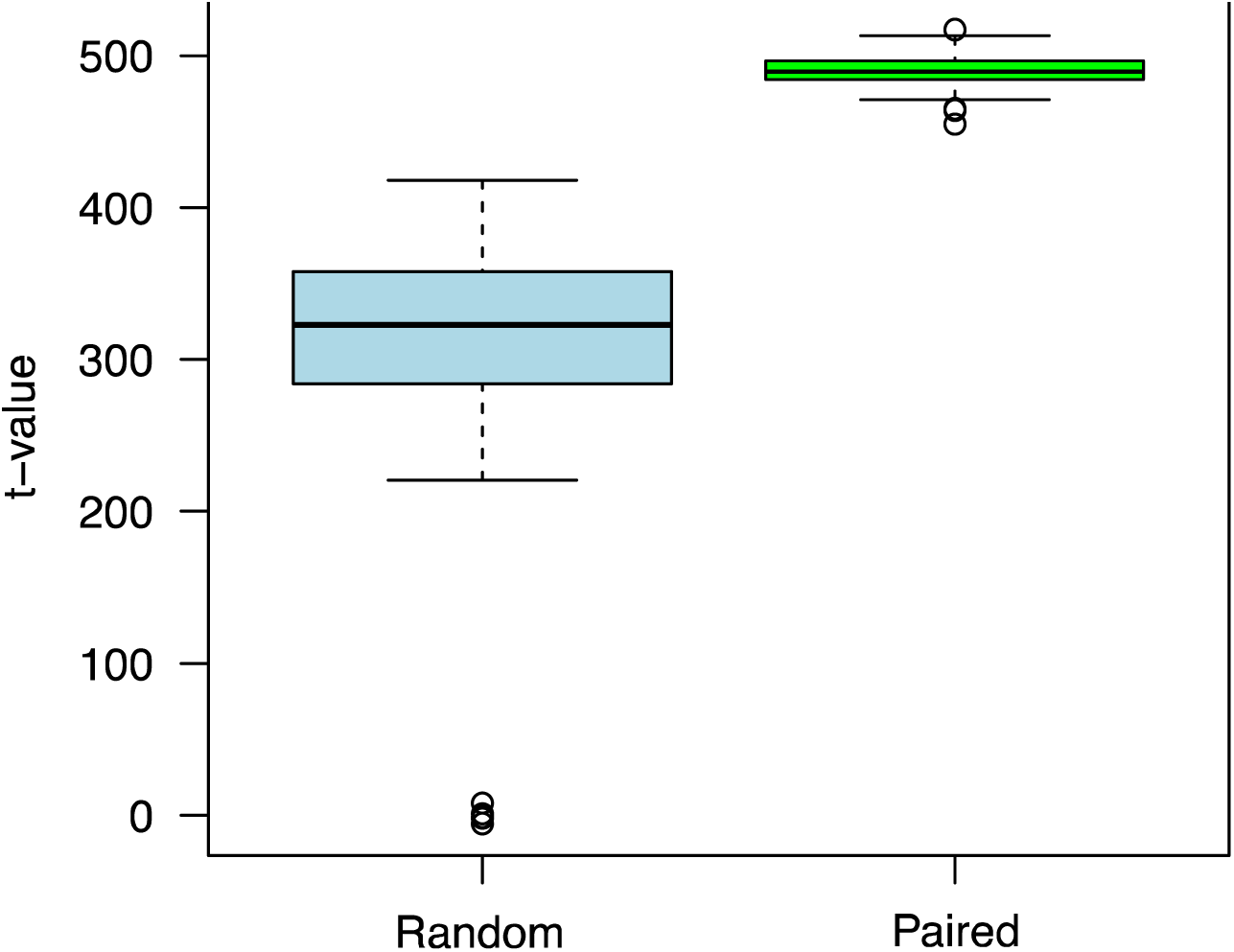
Summary data (boxplot) of model parameters (t-value) assessing facilitation effects in a spatial grid (abundance gradient codispersion). Plant species were sampled either randomly (blue) or paired (green). Lines represent Median, Q1 and Q3 as well as outlier range.

### Field results

At the Spanish site, there were highly significant beneficiary and stress level effects due to the lower cover of *Euphorbia* than of *Astydamia* and an overall decreasing cover with increasing stress, respectively (Fig. 6, Table S3). However, there was a significant stress by neighbour by beneficiary interaction because the decreasing cover with increasing stress was stronger for *Astydamia* in the open than below *Euphorbia* nurses, while there was a tendency for *Euphorbia* seedlings to increase in cover in the open but not below the nurse shrub. There were no significant effects of the method either as a single factor or in interaction with other factors.

**Fig. 6.**
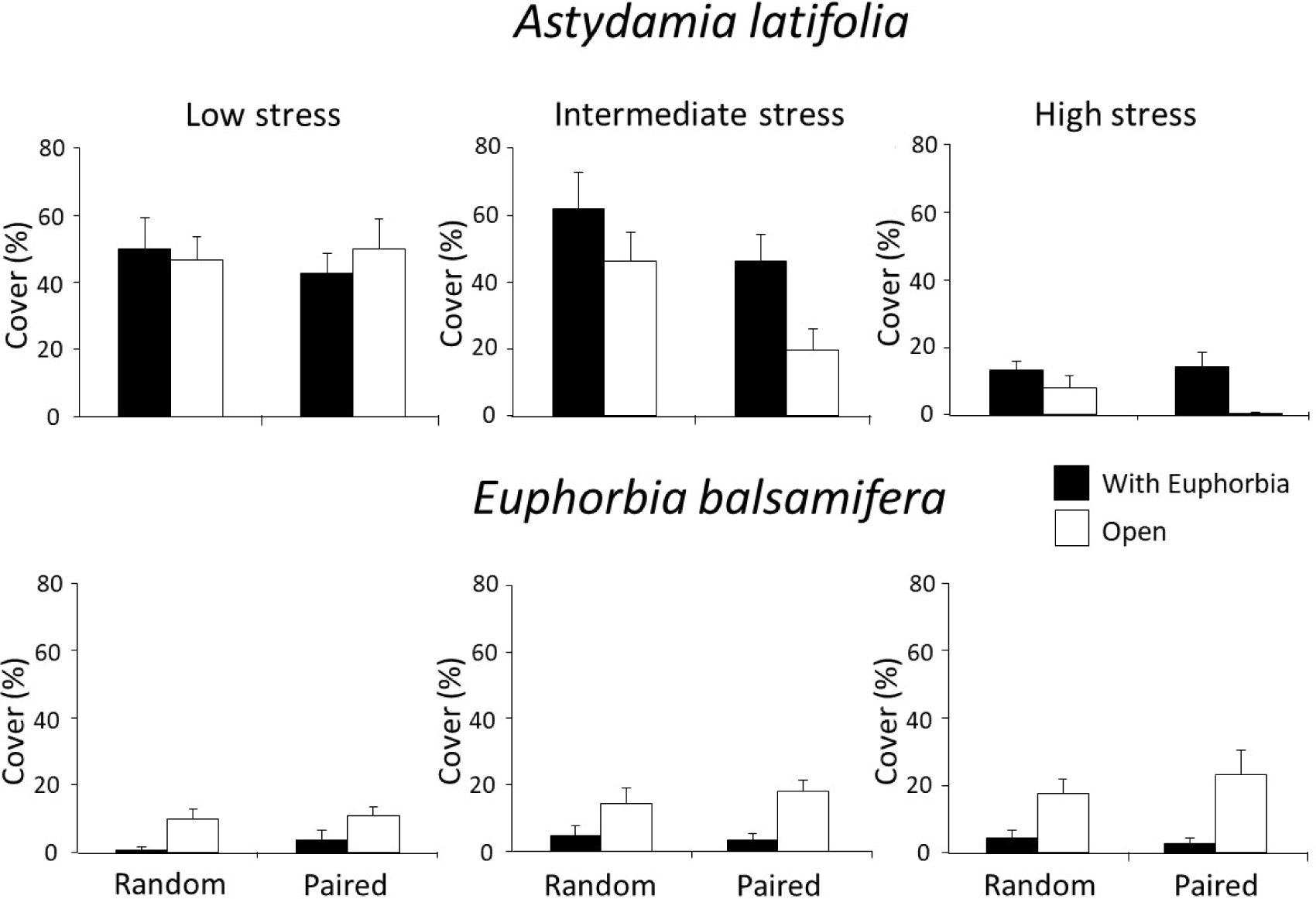
Means (± SE) of percent cover of *Astydamia* (upper panels) and *Euphorbia* (lower panels) beneficiaries with *Euphorbia* nurses and in open plots measured with the random (n = 14-46) and paired (n = 30) sampling methods, at three levels of stress (low: left panels, intermediate middle panels and high: right panels) at the Spanish site. See Table S1 for the complete statistical analyses.

At the Italian site, there were highly significant canopy position, neighbour and beneficiary effects, due to lower cover at the sunniest (south) canopy position, in the open and for *Cynara* than at the other two canopy positions, below the tree canopy and for *Smyrnium* (Fig. 7, Table S4). However, there was a highly significant canopy position by neighbour by beneficiary interaction because the lower abundance at the south position was observed for *Smyrnium* below the tree canopy while its abundance in the open was always very low and there were no differences among canopy positions for *Cynara* both under the tree canopy and in the open. There was also a significant method by beneficiary interaction because the abundance of *Smyrnium* was slightly higher with the paired than random method and the reverse was observed for *Cynara*. Finally, there was a weakly significant canopy position by method by neighbour interaction.

**Fig. 7.**
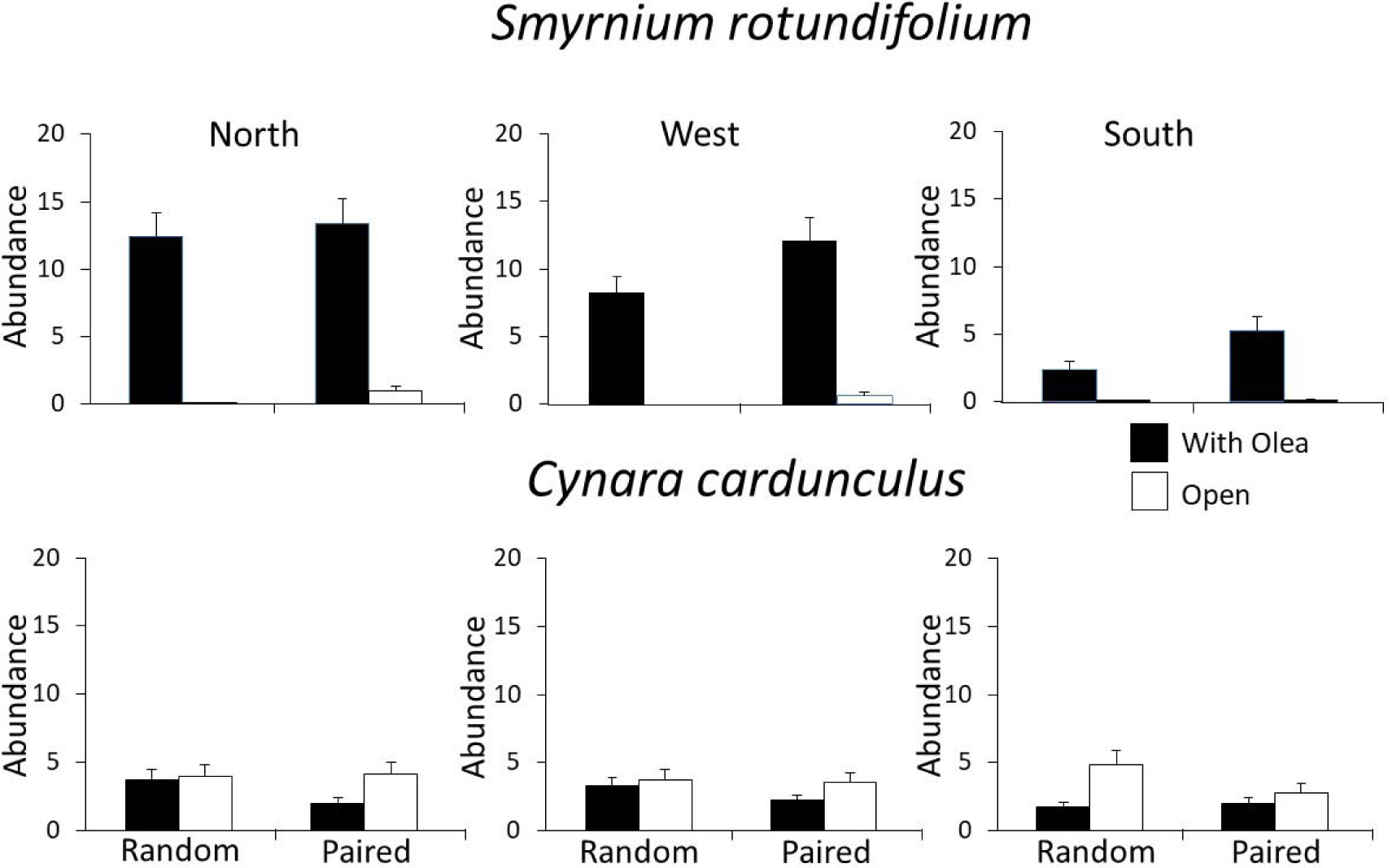
Mean (± SE, n = 30) abundance of *Smyrnium* (upper panels) and *Cynara* (lower panels) targets below *Olea* canopy and in the open measured with the random and paired sampling methods, at three canopy positions (North: left panels, West middle panels and South: right panels) at the Italian site. See Table S2 for the complete statistical analyses.

At the Japanese site, the performance of the two sampling techniques was very similar at the community level, with both methods showing more species associated with *Diapensia* cushions than colonizing open areas (Fig. 8). Additionally, there was no significant method x neighbour interaction and, thus, no difference in facilitation measured by the two methods (Table S5). For an abundant beneficiary species (*Salix nummularia*), frequency of occurrence was higher within *Diapensia* cushions than in the open with both methods (highly significant neighbour effect, Table S5). Additionally, frequency of occurrence was higher with the paired than random method (significant method effect, Table S5), due to the larger plot size used in the former than the latter (mean plot size of ca. 1000 cm^2^, vs. 100 cm^2^, respectively).

**Fig. 8.**
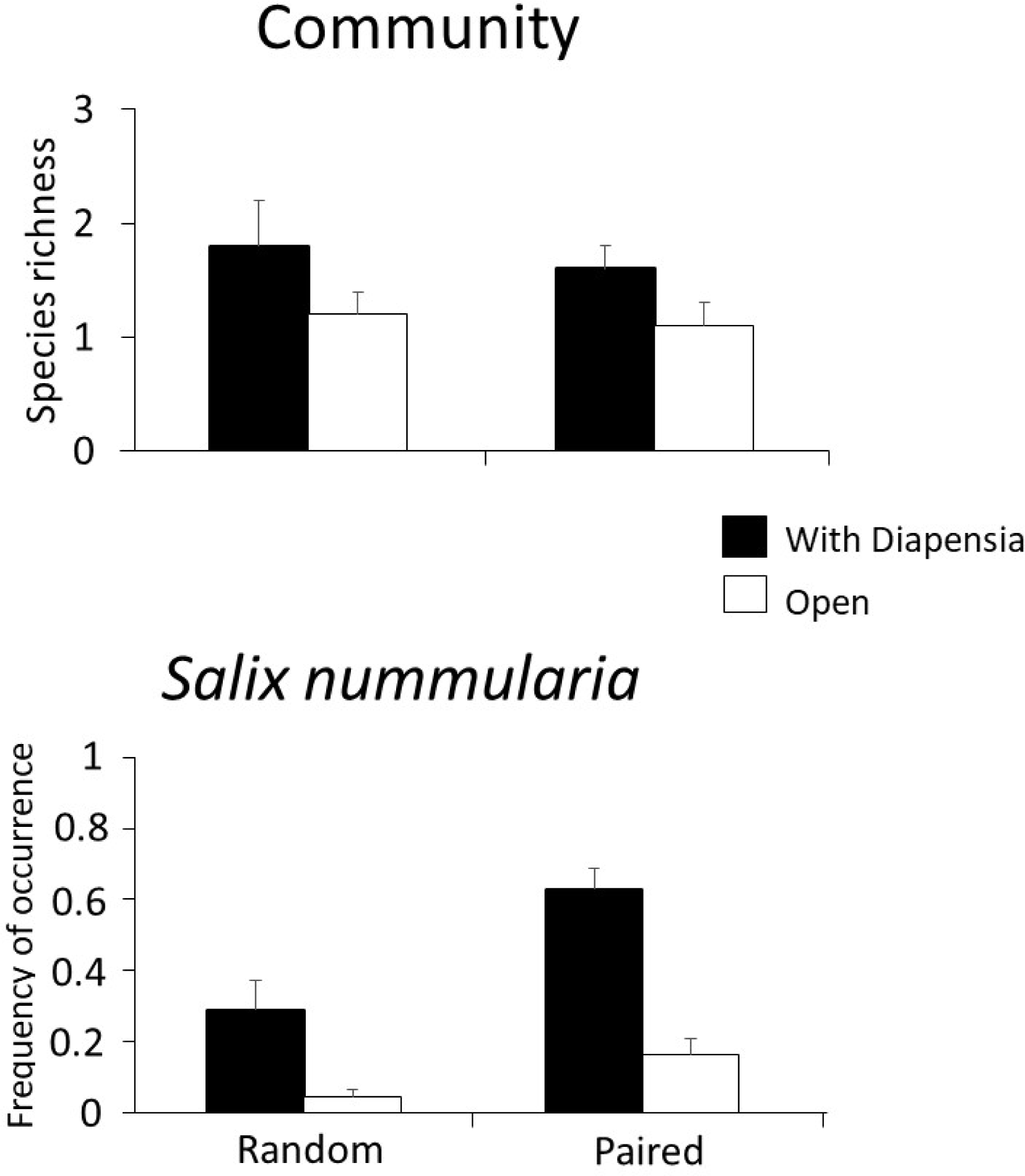
Mean (± SE, n = 70) species richness of the community (upper panel) and frequency of occurrence of *Salix* (lower panel) within the cushion canopy and in the open measured with the random and paired sampling methods at the Japanese site. See Table S3 for the complete statistical analyses.

## DISCUSSION

Our simulation results and field surveys indicate that the two methods of sampling pairwise or randomly provide similar outcomes. Notably, the methods were robust against local environmental heterogeneity. Yet, our simulation results showed that increasing beneficiary preference for the nurse tended to overestimate positive spatial associations when using the paired sampling method as compared to the random sampling method. Nevertheless, all other community parameters had no effect on differences between the methods in the observed non-random spatial associations. Additionally, spatial associations measured in the three systems were strongly influenced by species preferences and relative environmental stress. However, neither species nor stress effects led to differences in spatial associations measured with the two methods. Overall, these results show that the paired sampling method is as accurate as the random sampling method and, thus, can be used for assessing long-term facilitation in a wide range of environmental and community conditions, a task that cannot be accomplished with the removal method.

### The validity of the paired sampling method

The results of our simulated plant community indicate that the two sampling methods produce statistically indistinguishable estimates of facilitation effects. Yet, when considering facilitation as the absolute dependency of a beneficiary on a nurse plant (i.e., model parameter *b*_*1*_), nurse abundance and nurse environmental preferences, along with beneficiary preference for the nurse, influenced estimates of nurse effects detected by random sampling. Instead, only beneficiary preference for nurse influenced nurse effect estimates detected by the paired sampling method, as one would correctly expect. This indicates that random sampling is more sensitive to changing local conditions, but less accurate and robust than paired sampling in estimating species dependencies. When considering relative differences in between the two methods, such differences were negligible overall. Thus, within an overall picture of minimal influence of sampling method on estimating facilitation, the paired sampling does a better job in excluding the effects of local conditions and ultimately estimating the dependency of beneficiary plants on nurse species.

Some discrepancies arise between ours and previous simulations (Steinbauer et al. 2016). A possible reason is that what was previously defined as environmental heterogeneity was actually habitat sharing. In that case, all plants respond to environmental conditions the same way as they all show the same preferences for a ‘better’, less stressful environment.

However, this assumption is not very realistic since plants respond in different ways to environmental conditions (Liancourt et al. 2005b). In our model, we included a species-specific response to environmental conditions with contrasting preferences between plants for the environment. This way, we could separate the environmental heterogeneity component from the habitat-sharing component. Another likely explanation resides in the differences between modeling parameters. While the entire landscape is covered by plants and each cell has on average more than two plants in previous models (Steinbauer et al. 2016), we considered a more realistic stressful environment where plant cover does not exceed 50% of the landscape. Previous models also use many species, their simulation is based on individuals, and they consider species richness as the response variable (though without a Poisson or Negative Binomial distribution). On the other hand, our model considers only one beneficiary species, and species occurrence is used as the response variable (with a binomial distribution).

At the Japanese site we did not find any difference in spatial associations measured with the two methods. However, at this site within-community environmental pre-existent heterogeneity was low and we might expect no or little differences between methods. In contrast, at the Spanish field site within-community environmental heterogeneity is very high and it would be here where results of the paired sampling method were most likely to be affected by habitat-sharing effects. However, results showed no effects of the method. This shows that in sites with high microenvironmental heterogeneity, so long as the open and with-neighbour plots are sampled conservatively (i.e., not increasing the probability of occurrence of positive associations by sampling open plots in obviously more stressful microhabitats than with-neighbour plots) there is no impact of the method on the assessment of spatial association.

Finally, the Italian site contained low within-community environmental heterogeneity due to the flat topography. There, we found a weak but significant neighbour by canopy position by method interaction. However, since the effect was only weakly significant and occurred at the site with the lowest within-community environmental heterogeneity, processes other than habitat-sharing effects are likely to drive this effect.

### Species preferences and stress level influences

The only parameter that influenced spatial associations in our modeling study was beneficiary preference for the nurse. There was a tendency for more positive spatial associations with the paired than random sampling method when increasing beneficiary preference for the nurse. We also found strong differences in spatial associations with the dominants depending on the target species in both the Spanish and Italian systems. Additionally, we found strong variation in spatial associations with increasing stress in the Spanish system for *Astydamia*, the least stress-tolerant beneficiary species at this very dry site.

Plant communities include species from contrasting functional strategies with different responses to the abiotic environment and the effects of neighbours (Michalet et al. 2015b). At the Spanish site, *Euphorbia* increased in dominance with increasing stress, but the converse was observed for *Astydamia*, which is consistent with the former being present in drier conditions than the latter. In agreement with the functional tradeoffs described in several studies between physical stress- and shade-tolerance (Liancourt et al. 2005b, Nemer et al. 2021), *Euphorbia* was outcompeted at the three stress levels, whereas *Astydamia* was increasingly facilitated with increasing stress. At the Italian site, *Smyrnium* was strongly facilitated by *Olea*, whereas *Cynara* was more abundant in the open than below the trees, thus, highlighting that the former was less tolerant to high irradiance and the latter more negatively affected by shade. The former -less stress-tolerant -species showed higher variation in the effect of neighbours with canopy position than the shade-avoiding species, with higher facilitation found in the North position where light interception was the highest.

In conclusion, our results showed that the paired sampling method is robust enough for assessing spatial associations in a wide range of environmental and community conditions. Combining the paired sampling with other approaches may help to tease out confounding, habitat-sharing effects and identify the mechanisms underlying biotic interactions. This is important given the need to use the paired sampling approach in association with the removal method to enable disentangling of short-from long-term effects of neighbours (Michalet 2006; Schöb et al. 2012; Michalet et al. 2015a; Chaieb et al., 2021).

## Supporting information

Supplementary Information Talbe

## ACKNOWLEDGMENTS

GL was supported by the Swiss National Science Foundation (PZ00P3_202127).

## DATA AVAILABILITY STATEMENT

Data and code will be archived in Dryad.

