## Supplementary Information Code for "Assessing the accuracy of paired and random sampling for quantifying plant–plant interactions in natural communities"

Supplementary material - simulation R script

### Introduction

This Supplemetary Material reports the code to reproduce the modelling (simulation) and related figures carried out. The scope of this tutorial is to increase its reproducibility, clarity, transparency and dissemination. An HPC cluster is reccommended for using this tutorial effectively. Data analysis was done using R version 4.0.2. This document was compiled with the 'rmarkdown package, version Version 1.2.5033. This tutorial is licensed under CC BY-NC-ND 4.0, which means you are free to share, copy and redistribute this tutorial in any medium or format under the terms of attribution of appropriate credit, non-commercial purposes and no derivatives.

#### Software preparation

Install R (https://www.r-project.org) (R Core Team 2020) if you do not have it already. Then, install and load the following packages.

```
library(parallel)
library(foreach)
library(doParallel)
library(MASS)
library(car)
library(effects)
library(spatstat)
library(geoR)
library(fields)
library(SpatialPack)
library(grid)
library(raster)
library(gstat)
```

### Simulating plant communities

The plant community simulation consisted in two modelling approaches. We first assumed an homogeneous space, then we considered a gradient over a spatial grid.

#### Homogeneous space

We create a plant community composed of a nurse plant species $\alpha$ and a beneficiary plant species $\beta$. The community is assembled at different levels of environmental heterogeneity, environmental severity, plant abundance, and plant preferences. Two levels of sampling effort are considered (i.e., 5% and 10%) for both the random or pairwise methods.

```
# environmental homogeneity
env_h = gl(2, 1, label=c('homo', 'hetero'))
# environmental quality
```

```
env_q = c(0.2, 0.9)
# nurse abundance
nur_a = seq(0.05, 0.3, length.out = 4)
# beneficiary abundance
ben_a = seq(0.01, 0.2, length.out = 4)
# nurse preference
nur_p = seq(0.2, 1, length.out = 4)
# beneficiary preference to environment
ben_p = seq(0.2, 1, length.out = 4)
# beneficiary preference to nurses
ben_n = seq(0, 1, length.out = 4)
# sampling effort
sampl = c(0.025, 0.05)*250000
# data frame for storing results
study_df = expand.grid(env_h,nur_a, ben_a, nur_p, ben_p, ben_n,sampl)
colnames(study_df) = c('env_h','nur_a','ben_a','nur_p','ben_p','ben_n','sampl')
casi = nrow(study_df)
study_df$random = NA
study_df$paired = NA
## create list for storing data
study_list = rep(list(rep(list(NA),3)),casi)
for(i in 1:casi){
print(c(i, casi-i))
##### create environment
if(study_df[i,1]=='homo') p_e = rep(0.1,250000)
else p_e = sample(c(0.1,0.9),250000, replace=T, prob = c(0.7,0.3))
##### create community
### nurse
## "niche matching" considers combination of env quality and nurse preference
nm = 1-abs(p_e-study_df[i,4])
## prob occurrence (prese abs) nurse
pa_nu = rbinom(250000, 1, sqrt(study_df[i,2]*nm))
### benef preference
## include nurse presence
nm = 1-abs(p_e-study_df[i,5])
np = 1-abs(pa_nu-study_df[i,6])
pa_be = rbinom(250000, 1, sqrt(sqrt(study_df[i,3]*nm)*np))
###### random sampling
sampl_r = sample(1:250000, study_df$sampl[i]*2)
prand = data.frame(nurse = pa_nu[sampl_r], benef = pa_be[sampl_r])
###### paired sampling
sampl_n = sample(which(pa_nu==1), study_df$sampl[i])
sampl_o = sample(which(pa_nu==0), study_df$sampl[i])
ppair = data.frame(nurse = c(pa_nu[sampl_n],pa_nu[sampl_o]),
```

```
                        benef = c(pa_be[sampl_n],pa_be[sampl_o]))
### store data
study_list[[i]][[1]] = cbind(pa_nu, pa_be)
study_list[[i]][[2]] = prand
study_list[[i]][[3]] = ppair
}
```

##### Statistical analysis of simulated plant communities

We addressed the influence of sampling methods on estimated facilitation effects. Facilitation effects were calculated as the dependency of beneficiary species on nurse species using a generalized linear model.

```
## stima b_1 parms
random_b <- foreach(i=1:casi, .inorder = TRUE, .combine=c) %dopar% {
    glm1 <- glm(benef~nurse, data=study_list[[i]][[2]],
                family="binomial")
    summary(glm1)$coefficients[2,1]
}

stopImplicitCluster()

paired_b <- foreach(i=1:casi, .inorder = TRUE, .combine=c) %dopar% {
    glm1 <- glm(benef~nurse, data=study_list[[i]][[3]],
                family="binomial")
    summary(glm1)$coefficients[2,1]
}

paired_b = c(paired1, paired2, paired3, paired4)

stopImplicitCluster()

study_df$random_b = random_b
study_df$paired_b = paired_b

## z-value
random <- foreach(i=1:casi, .inorder = TRUE, .combine=c) %dopar% {
    glm1 <- glm(benef~nurse, data=study_list[[i]][[2]],
                family="binomial")
    summary(glm1)$coefficients[2,3]
}

stopImplicitCluster()

paired <- foreach(i=1:casi, .inorder = TRUE, .combine=c) %dopar% {
```

```
    glm1 <- glm(benef~nurse, data=study_list[[i]][[3]],
                family="binomial")
    summary(glm1)$coefficients[2,3]
}

stopImplicitCluster()

study_df$random = random
study_df$paired = paired
```

Then, we looked at the impact of abiotic and biotic factors on both raw and normalized (i.e., $z$-score) $b_1$ parameters for each sampling method separately prior to addressing the relative differences $\delta$ in facilitation effects between pairwise and random sampling.

##### Estimate $b_1$ as response

```
cor.test(random_b,paired_b, "pearson", alternative = "two.sided")
```

```
##
##  Pearson's product-moment correlation
##
## data:  random_b and paired_b
## t = 1234.9, df = 4094, p-value < 2.2e-16
## alternative hypothesis: true correlation is not equal to 0
## 95 percent confidence interval:
##  0.9985757 0.9987399
## sample estimates:
##       cor
## 0.9986603
```

```
#######

mod_random_b = lm(random_b~ env_h+nur_a+ben_a+nur_p+ben_p+ben_n+sampl,
                  data = study_df)
Anova(mod_random_b)
```

```
## Anova Table (Type II tests)
##
## Response: random_b
##          Sum Sq   Df    F value    Pr(>F)
## env_h         5    1     0.3149 0.5747233
## nur_a       194    1    11.4376 0.0007265 ***
## ben_a         0    1     0.0027 0.9582987
## nur_p       140    1     8.2812 0.0040265 **
## ben_p         1    1     0.0349 0.8518440
```

```
## ben_n     721082    1 42521.7971 < 2.2e-16 ***
## sampl          0    1     0.0003 0.9863357
## Residuals  69324 4088
## ---
## Signif. codes:  0 '***' 0.001 '**' 0.01 '*' 0.05 '.' 0.1 ' ' 1
```

```
summary(mod_random_b)
```

```
##
## Call:
## lm(formula = random_b ~ env_h + nur_a + ben_a + nur_p + ben_p +
##     ben_n + sampl, data = study_df)
##
## Residuals:
##     Min      1Q  Median      3Q     Max
## -6.7320 -3.4089  0.6113  4.1476  5.6518
##
## Coefficients:
##                  Estimate Std. Error t value Pr(>|t|)
## (Intercept) -1.722e+01  3.320e-01 -51.877  < 2e-16 ***
## env_hhetero -7.221e-02  1.287e-01  -0.561 0.574723
## nur_a       -2.336e+00  6.906e-01  -3.382 0.000726 ***
## ben_a        4.752e-02  9.087e-01   0.052 0.958299
## nur_p        6.211e-01  2.158e-01   2.878 0.004026 **
## ben_p        4.031e-02  2.158e-01   0.187 0.851844
## ben_n        3.560e+01  1.727e-01 206.208  < 2e-16 ***
## sampl       -3.527e-07  2.059e-05  -0.017 0.986336
## ---
## Signif. codes:  0 '***' 0.001 '**' 0.01 '*' 0.05 '.' 0.1 ' ' 1
##
## Residual standard error: 4.118 on 4088 degrees of freedom
## Multiple R-squared:  0.9123, Adjusted R-squared:  0.9122
## F-statistic:  6077 on 7 and 4088 DF,  p-value: < 2.2e-16
```

```
mod_paired_b = lm(paired_b~ env_h+nur_a+ben_a+nur_p+ben_p+ben_n+sampl,
                  data = study_df)
Anova(mod_paired_b)
```

```
## Anova Table (Type II tests)
##
## Response: paired_b
##           Sum Sq   Df    F value Pr(>F)
## env_h          0    1     0.0013 0.9710
## nur_a          0    1     0.0000 0.9986
## ben_a          0    1     0.0035 0.9526
## nur_p          0    1     0.0167 0.8971
```

```
## ben_p          0    1     0.0250 0.8743
## ben_n     699933    1 43704.4304 <2e-16 ***
## sampl          0    1     0.0000 0.9951
## Residuals  65470 4088
## ---
## Signif. codes:  0 '***' 0.001 '**' 0.01 '*' 0.05 '.' 0.1 ' ' 1
```

```
summary(mod_paired_b)
```

```
##
## Call:
## lm(formula = paired_b ~ env_h + nur_a + ben_a + nur_p + ben_p +
##     ben_n + sampl, data = study_df)
##
## Residuals:
##     Min      1Q  Median      3Q     Max
## -5.6215 -3.2609 -0.0138  3.2936  5.6887
##
## Coefficients:
##               Estimate Std. Error t value Pr(>|t|)
## (Intercept) -1.758e+01  3.227e-01 -54.488   <2e-16 ***
## env_hhetero  4.543e-03  1.251e-01   0.036    0.971
## nur_a        1.164e-03  6.711e-01   0.002    0.999
## ben_a        5.250e-02  8.831e-01   0.059    0.953
## nur_p        2.714e-02  2.097e-01   0.129    0.897
## ben_p        3.318e-02  2.097e-01   0.158    0.874
## ben_n        3.508e+01  1.678e-01 209.056   <2e-16 ***
## sampl        1.233e-07  2.001e-05   0.006    0.995
## ---
## Signif. codes:  0 '***' 0.001 '**' 0.01 '*' 0.05 '.' 0.1 ' ' 1
##
## Residual standard error: 4.002 on 4088 degrees of freedom
## Multiple R-squared:  0.9145, Adjusted R-squared:  0.9143
## F-statistic:  6243 on 7 and 4088 DF,  p-value: < 2.2e-16
```

Results indicate that the two methods produce highly correlated facilitation effects.

Nurse abundance (`nur_a`), nurse environmental preference (`nur_p`), and beneficiary preference for nurse (`ben_a`) influenced the nurse effect.

Instead, only beneficiary preference for nurse (`ben_a`) influenced the nurse effect in paired sampling.

In summary, paired sampling is more robust than random one.

##### Figures

```
par(mfrow=c(2,4), mar=c(4,4,1.2,1.2))
## A
plot(random_b,paired_b, las=1, col="black",
xlab="random [beta]",
ylab = "paired [beta]",
xlim = range(c(paired_b,random_b)),
ylim = range(c(paired_b,random_b)))
title('A', adj = 0, line = 0)
abline(c(0, 1))
abline(h=1.96, lty=2)
abline(h=-1.96, lty=2)
abline(v=1.96, lty=2)
abline(v=-1.96, lty=2)
legend("topleft", legend=c("cor=0.998", "p < 0.001"),
       bty="n", inset=0, cex=.75)
## B
plot(0,0,type="n",xlim=c(1,2), ylim=range(c(paired_b,random_b)),
     xaxt="n",xlab="Environment",
     ylab="Nurse effect [beta]",yaxt="n")
title('B', adj = 0, line = 0)
axis(2,at=c(-20,-10,0,10,20), las=1)
axis(1,at=c(1,2), labels=c('homogeneous','heterogeneous'), las=1)
points(rep(1,casi/2), random_b[which(study_df$env_h=='homo')],
       pch = 21, col = 'blue3')
points(rep(1.8,casi/2), random_b[which(study_df$env_h=='hetero')],
       pch = 21, col = 'blue3')
points(rep(1.2,casi/2), paired_b[which(study_df$env_h=='homo')],
       pch = 21, col = 'red')
points(rep(2,casi/2), paired_b[which(study_df$env_h=='hetero')],
       pch = 21, col = 'red')
abline(coef(mod_random_b)[1], coef(mod_random_b)[2], col='blue3')
abline(coef(mod_paired_b)[1], coef(mod_paired_b)[2], col='red')
## C
plot(0,0,type="n",xlim=c(0.05,0.31), ylim=range(c(paired_b,random_b)),
     xaxt="n",xlab="Nurse abundance [%]",
     ylab="Nurse effect [beta]",yaxt="n")
title('C', adj = 0, line = 0)
axis(2,at=c(-20,-10,0,10,20), las=1)
axis(1,at=c(0.05, 0.1, 0.2, 0.3), las=1)
points(study_df[,2], random_b, pch = 21, col = 'blue3')
points(study_df[,2]+0.01, paired_b, pch = 21, col = 'red')
abline(coef(mod_random_b)[1], coef(mod_random_b)[3], col='blue3')
```

```
abline(coef(mod_paired_b)[1], coef(mod_paired_b)[3], col='red')
## D
plot(0,0,type="n",xlim=c(0.01,0.21), ylim=range(c(paired_b,random_b)),
     xaxt="n",xlab="Beneficiary abundance [%]",
     ylab="Nurse effect [beta]",yaxt="n")
title('D', adj = 0, line = 0)
axis(2,at=c(-20,-10,0,10,20), las=1)
axis(1,at=c(0.01, 0.05, 0.10, 0.15, 0.20), las=1)
points(study_df[,3], random_b, pch = 21, col = 'blue3')
points(study_df[,3]+0.01, paired_b, pch = 21, col = 'red')
abline(coef(mod_random_b)[1], coef(mod_random_b)[4], col='blue3')
abline(coef(mod_paired_b)[1], coef(mod_paired_b)[4], col='red')
## E
plot(0,0,type="n",xlim=c(0.2,1.03), ylim=range(c(paired_b,random_b)),
     xaxt="n",xlab="Nurse environmental preference [%]",
     ylab="Nurse effect [beta]",yaxt="n")
title('E', adj = 0, line = 0)
axis(2,at=c(-20,-10,0,10,20), las=1)
axis(1,at=c(0.2, 0.4, 0.6, 0.8, 1.0), las=1)
points(study_df[,4], random_b, pch = 21, col = 'blue3')
points(study_df[,4]+0.03, paired_b, pch = 21, col = 'red')
abline(coef(mod_random_b)[1], coef(mod_random_b)[5], col='blue3')
abline(coef(mod_paired_b)[1], coef(mod_paired_b)[5], col='red')
## F
plot(0,0,type="n",xlim=c(0.2,1.03), ylim=range(c(paired_b,random_b)),
     xaxt="n",xlab="Beneficiary environmental preference [%]",
     ylab="Nurse effect [beta]",yaxt="n")
title('F', adj = 0, line = 0)
axis(2,at=c(-20,-10,0,10,20), las=1)
axis(1,at=c(0.2, 0.4, 0.6, 0.8, 1.0), las=1)
points(study_df[,5], random_b, pch = 21, col = 'blue3')
points(study_df[,5]+0.03, paired_b, pch = 21, col = 'red')
abline(coef(mod_random_b)[1], coef(mod_random_b)[6], col='blue3')
abline(coef(mod_paired_b)[1], coef(mod_paired_b)[6], col='red')
## G
plot(0,0,type="n",xlim=c(0,1.03), ylim=range(c(paired_b,random_b)),
     xaxt="n",xlab="Beneficiary preference for nurse [%]",
     ylab="Nurse effect [beta]",yaxt="n")
title('G', adj = 0, line = 0)
axis(2,at=c(-20,-10,0,10,20), las=1)
axis(1,at=c(0, 0.2, 0.4, 0.6, 0.8, 1.0), las=1)
points(study_df[,6], random_b, pch = 21, col = 'blue3')
points(study_df[,6]+0.03, paired_b, pch = 21, col = 'red')
abline(coef(mod_random_b)[1], coef(mod_random_b)[7], col='blue3')
```

```r
abline(coef(mod_paired_b)[1], coef(mod_paired_b)[7], col='red')
legend("topleft", legend=c("random", "paired"), fill=c('blue3','red'),
       bty="n", inset=0, cex=.75)
## H
plot(0,0,type="n",xlim=c(6250, 12700), ylim=range(c(paired_b,random_b)),
     xaxt="n",xlab="Sampling effort [%]",
     ylab="Nurse effect [beta]",yaxt="n")
title('H', adj = 0, line = 0)
axis(2,at=c(-20,-10,0,10,20), las=1)
axis(1,at=c(6250, 12500), labels=c(5, 10), las=1)
points(study_df[,7], random_b, pch = 21, col = 'blue3')
points(study_df[,7]+250, paired_b, pch = 21, col = 'red')
abline(coef(mod_random_b)[1], coef(mod_random_b)[8], col='blue3')
abline(coef(mod_paired_b)[1], coef(mod_paired_b)[8], col='red')
```

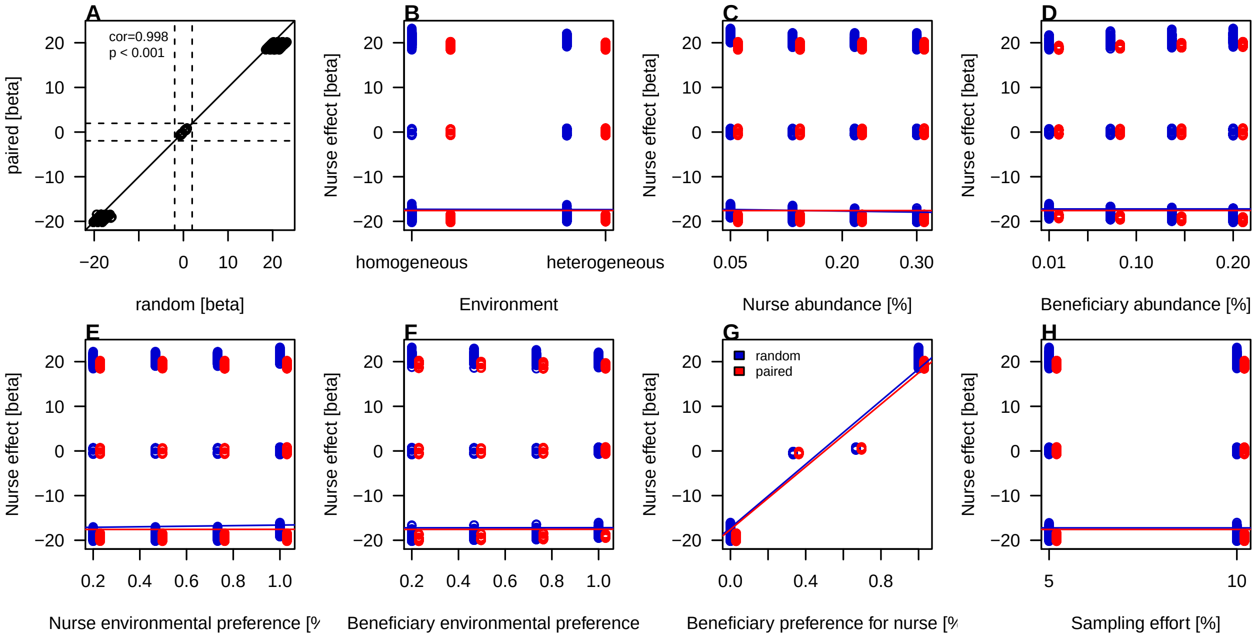

##### Normalized *z*-score as response

```r
cor.test(random,paired, "pearson", alternative = "two.sided")
```

```
##
##  Pearson's product-moment correlation
##
## data:  random and paired
## t = 372.51, df = 4094, p-value < 2.2e-16
## alternative hypothesis: true correlation is not equal to 0
## 95 percent confidence interval:
##  0.9846619 0.9864189
```

```
## sample estimates:
##       cor
## 0.9855669
```

```
#####

mod_random_z = lm(random~ env_h+nur_a+ben_a+nur_p+ben_p+ben_n+sampl,
                  data = study_df)
Anova(mod_random_z)
```

```
## Anova Table (Type II tests)
##
## Response: random
##           Sum Sq   Df  F value Pr(>F)
## env_h         14    1   0.1693 0.6808
## nur_a          4    1   0.0473 0.8279
## ben_a          1    1   0.0175 0.8947
## nur_p        192    1   2.2485 0.1338
## ben_p        163    1   1.9105 0.1670
## ben_n      36607    1 428.8067 <2e-16 ***
## sampl          4    1   0.0468 0.8287
## Residuals 348989 4088
## ---
## Signif. codes:  0 '***' 0.001 '**' 0.01 '*' 0.05 '.' 0.1 ' ' 1
```

```
summary(mod_random_z)
```

```
##
## Call:
## lm(formula = random ~ env_h + nur_a + ben_a + nur_p + ben_p +
##     ben_n + sampl, data = study_df)
##
## Residuals:
##      Min       1Q   Median       3Q      Max
## -24.6787  -4.5734  -0.3177   4.2807  25.4549
##
## Coefficients:
##                 Estimate Std. Error t value Pr(>|t|)
## (Intercept) -4.767e+00  7.450e-01  -6.399 1.75e-10 ***
## env_hhetero  1.188e-01  2.887e-01   0.411    0.681
## nur_a       -3.370e-01  1.550e+00  -0.217    0.828
## ben_a       -2.700e-01  2.039e+00  -0.132    0.895
## nur_p        7.261e-01  4.842e-01   1.500    0.134
## ben_p        6.693e-01  4.842e-01   1.382    0.167
## ben_n        8.022e+00  3.874e-01  20.708  < 2e-16 ***
## sampl        9.997e-06  4.620e-05   0.216    0.829
```

```
## ---
## Signif. codes:  0 '***' 0.001 '**' 0.01 '*' 0.05 '.' 0.1 ' ' 1
##
## Residual standard error: 9.24 on 4088 degrees of freedom
## Multiple R-squared:  0.09582,    Adjusted R-squared:  0.09428
## F-statistic: 61.89 on 7 and 4088 DF,  p-value: < 2.2e-16
```

```
mod_paired_z = lm(paired~ env_h+nur_a+ben_a+nur_p+ben_p+ben_n+sampl,
                  data = study_df)
Anova(mod_paired_z)
```

```
## Anova Table (Type II tests)
##
## Response: paired
##           Sum Sq   Df  F value Pr(>F)
## env_h         30    1   0.2589 0.6109
## nur_a          0    1   0.0003 0.9856
## ben_a          2    1   0.0138 0.9066
## nur_p        159    1   1.3965 0.2374
## ben_p        239    1   2.0976 0.1476
## ben_n      50295    1 440.6267 <2e-16 ***
## sampl          2    1   0.0159 0.8997
## Residuals 466619 4088
## ---
## Signif. codes:  0 '***' 0.001 '**' 0.01 '*' 0.05 '.' 0.1 ' ' 1
```

```
summary(mod_paired_z)
```

```
##
## Call:
## lm(formula = paired ~ env_h + nur_a + ben_a + nur_p + ben_p +
##     ben_n + sampl, data = study_df)
##
## Residuals:
##      Min       1Q   Median       3Q      Max
## -28.0986  -5.2760   0.4158   5.1346  28.9421
##
## Coefficients:
##               Estimate Std. Error t value Pr(>|t|)
## (Intercept) -5.667e+00  8.614e-01  -6.579 5.34e-11 ***
## env_hhetero  1.699e-01  3.339e-01   0.509    0.611
## nur_a       -3.245e-02  1.792e+00  -0.018    0.986
## ben_a        2.766e-01  2.358e+00   0.117    0.907
## nur_p        6.617e-01  5.599e-01   1.182    0.237
## ben_p        8.109e-01  5.599e-01   1.448    0.148
## ben_n        9.403e+00  4.479e-01  20.991  < 2e-16 ***
```

```
## sampl        6.731e-06  5.342e-05   0.126    0.900
## ---
## Signif. codes:  0 '***' 0.001 '**' 0.01 '*' 0.05 '.' 0.1 ' ' 1
##
## Residual standard error: 10.68 on 4088 degrees of freedom
## Multiple R-squared:  0.09805,    Adjusted R-squared:  0.09651
## F-statistic: 63.49 on 7 and 4088 DF,  p-value: < 2.2e-16
```

No acutal differences between the two methods as the two facilitation effects are highly correlated.

In 'relative' terms, only beneficiary preference for nurse influences nurse effect in both random and sampling methods.

So, no real differences emerge between two methods when using a standardized and normalized effect variable.

```
study_df2 = rbind(study_df,study_df)
study_df2$method = gl(2, 4096, 2* 4096, c('random','paired'))
study_df2$zval = c(study_df$random, study_df$paired)

mod_r1 = lm(zval ~ method*(env_h+nur_a+ben_a+nur_p+ben_p+ben_n+sampl),
            data = study_df2)
Anova(mod_r1)
```

```
## Anova Table (Type II tests)
##
## Response: zval
##               Sum Sq   Df  F value  Pr(>F)
## method             7    1   0.0692 0.79254
## env_h             43    1   0.4277 0.51312
## nur_a              2    1   0.0243 0.87608
## ben_a              0    1   0.0000 0.99830
## nur_p            351    1   3.5146 0.06087 .
## ben_p            399    1   3.9984 0.04558 *
## ben_n          86359    1 865.7010 < 2e-16 ***
## sampl              6    1   0.0561 0.81277
## method:env_h       1    1   0.0134 0.90783
## method:nur_a       2    1   0.0165 0.89772
## method:ben_a       3    1   0.0308 0.86078
## method:nur_p       1    1   0.0076 0.93064
## method:ben_p       4    1   0.0366 0.84828
## method:ben_n     542    1   5.4371 0.01974 *
## method:sampl       0    1   0.0021 0.96313
## Residuals     815608 8176
## ---
## Signif. codes:  0 '***' 0.001 '**' 0.01 '*' 0.05 '.' 0.1 ' ' 1
```

```r
summary(mod_r1)
```

```
##
## Call:
## lm(formula = zval ~ method * (env_h + nur_a + ben_a + nur_p +
##     ben_p + ben_n + sampl), data = study_df2)
##
## Residuals:
##      Min       1Q   Median       3Q      Max
## -28.0986  -5.1040  -0.1865   4.9792  28.9421
##
## Coefficients:
##                             Estimate Std. Error t value Pr(>|t|)
## (Intercept)               -4.767e+00  8.053e-01  -5.919 3.37e-09 ***
## methodpaired              -9.004e-01  1.139e+00  -0.791   0.4292
## env_hhetero                1.188e-01  3.121e-01   0.381   0.7035
## nur_a                     -3.370e-01  1.675e+00  -0.201   0.8406
## ben_a                     -2.700e-01  2.204e+00  -0.123   0.9025
## nur_p                      7.261e-01  5.234e-01   1.387   0.1654
## ben_p                      6.693e-01  5.234e-01   1.279   0.2011
## ben_n                      8.022e+00  4.188e-01  19.156  < 2e-16 ***
## sampl                      9.997e-06  4.994e-05   0.200   0.8413
## methodpaired:env_hhetero  5.111e-02  4.414e-01   0.116   0.9078
## methodpaired:nur_a         3.045e-01  2.369e+00   0.129   0.8977
## methodpaired:ben_a         5.466e-01  3.117e+00   0.175   0.8608
## methodpaired:nur_p        -6.443e-02  7.403e-01  -0.087   0.9306
## methodpaired:ben_p         1.416e-01  7.403e-01   0.191   0.8483
## methodpaired:ben_n         1.381e+00  5.922e-01   2.332   0.0197 *
## methodpaired:sampl        -3.265e-06  7.062e-05  -0.046   0.9631
## ---
## Signif. codes:  0 '***' 0.001 '**' 0.01 '*' 0.05 '.' 0.1 ' ' 1
##
## Residual standard error: 9.988 on 8176 degrees of freedom
## Multiple R-squared:  0.09711,    Adjusted R-squared:  0.09545
## F-statistic: 58.62 on 15 and 8176 DF,  p-value: < 2.2e-16
```

The following factors and sampling method influence variance of facilitation effect: nurse habitat preference (marginally), beneficiary habitat preference, and beneficiary affinity to nurse alone and depending on sampling method.

When considering parameter estimates, beneficiary affinity to nurse significantly increases facilitation effects quantification, both per se, as reasonably expected, as well as with higher impact using the pairwise method.

**Figures**

```
par(mfrow=c(2,4), mar=c(4,4,1.2,1.2))
## A
plot(random,paired, las=1, col="black",
xlab="random [z-score]",
ylab = "paired [z-score]",
xlim = range(c(paired,random)),
ylim = range(c(paired,random)))
title('A', adj = 0, line = 0)
abline(c(0, 1))
abline(h=1.96, lty=2)
abline(h=-1.96, lty=2)
abline(v=1.96, lty=2)
abline(v=-1.96, lty=2)
legend("topleft", legend=c("cor=0.998", "p < 0.001"),
       bty="n", inset=0, cex=.75)
## B
plot(0,0,type="n",xlim=c(1,2), ylim=range(c(paired,random)),
     xaxt="n",xlab="Environment",
     ylab="Nurse effect [z-score]",yaxt="n")
title('B', adj = 0, line = 0)
axis(2,at=c(-20,-10,0,10,20), las=1)
axis(1,at=c(1,2), labels=c('homogeneous','heterogeneous'), las=1)
points(rep(1,casi/2), random[which(study_df$env_h=='homo')],
       pch = 21, col = 'blue3')
points(rep(1.8,casi/2), random[which(study_df$env_h=='hetero')],
       pch = 21, col = 'blue3')
points(rep(1.2,casi/2), paired[which(study_df$env_h=='homo')],
       pch = 21, col = 'red')
points(rep(2,casi/2), paired[which(study_df$env_h=='hetero')],
       pch = 21, col = 'red')
abline(coef(mod_random_z)[1], coef(mod_random_z)[2], col='blue3')
abline(coef(mod_paired_z)[1], coef(mod_paired_z)[2], col='red')
## C
plot(0,0,type="n",xlim=c(0.05,0.31), ylim=range(c(paired,random)),
     xaxt="n",xlab="Nurse abundance [%]",
     ylab="Nurse effect [z-score]",yaxt="n")
title('C', adj = 0, line = 0)
axis(2,at=c(-20,-10,0,10,20), las=1)
axis(1,at=c(0.05, 0.1, 0.2, 0.3), las=1)
points(study_df[,2], random, pch = 21, col = 'blue3')
points(study_df[,2]+0.01, paired, pch = 21, col = 'red')
abline(coef(mod_random_z)[1], coef(mod_random_z)[3], col='blue3')
```

```r
abline(coef(mod_paired_z)[1], coef(mod_paired_z)[3], col='red')
## D
plot(0,0,type="n",xlim=c(0.01,0.21), ylim=range(c(paired,random)),
     xaxt="n",xlab="Beneficiary abundance [%]",
     ylab="Nurse effect [z-score]",yaxt="n")
title('D', adj = 0, line = 0)
axis(2,at=c(-20,-10,0,10,20), las=1)
axis(1,at=c(0.01, 0.05, 0.10, 0.15, 0.20), las=1)
points(study_df[,3], random, pch = 21, col = 'blue3')
points(study_df[,3]+0.01, paired, pch = 21, col = 'red')
abline(coef(mod_random_z)[1], coef(mod_random_z)[4], col='blue3')
abline(coef(mod_paired_z)[1], coef(mod_paired_z)[4], col='red')
## E
plot(0,0,type="n",xlim=c(0.2,1.03), ylim=range(c(paired,random)),
     xaxt="n",xlab="Nurse environmental preference [%]",
     ylab="Nurse effect [z-score]",yaxt="n")
title('E', adj = 0, line = 0)
axis(2,at=c(-20,-10,0,10,20), las=1)
axis(1,at=c(0.2, 0.4, 0.6, 0.8, 1.0), las=1)
points(study_df[,4], random, pch = 21, col = 'blue3')
points(study_df[,4]+0.03, paired, pch = 21, col = 'red')
abline(coef(mod_random_z)[1], coef(mod_random_z)[5], col='blue3')
abline(coef(mod_paired_z)[1], coef(mod_paired_z)[5], col='red')
## F
plot(0,0,type="n",xlim=c(0.2,1.03), ylim=range(c(paired,random)),
     xaxt="n",xlab="Beneficiary environmental preference [%]",
     ylab="Nurse effect [z-score]",yaxt="n")
title('F', adj = 0, line = 0)
axis(2,at=c(-20,-10,0,10,20), las=1)
axis(1,at=c(0.2, 0.4, 0.6, 0.8, 1.0), las=1)
points(study_df[,5], random, pch = 21, col = 'blue3')
points(study_df[,5]+0.03, paired, pch = 21, col = 'red')
abline(coef(mod_random_z)[1], coef(mod_random_z)[6], col='blue3')
abline(coef(mod_paired_z)[1], coef(mod_paired_z)[6], col='red')
## G
plot(0,0,type="n",xlim=c(0,1.03), ylim=range(c(paired,random)),
     xaxt="n",xlab="Beneficiary preference for nurse [%]",
     ylab="Nurse effect [z-score]",yaxt="n")
title('G', adj = 0, line = 0)
axis(2,at=c(-20,-10,0,10,20), las=1)
axis(1,at=c(0, 0.2, 0.4, 0.6, 0.8, 1.0), las=1)
points(study_df[,6], random, pch = 21, col = 'blue3')
points(study_df[,6]+0.03, paired, pch = 21, col = 'red')
abline(coef(mod_random_z)[1], coef(mod_random_z)[7], col='blue3')
```

```
abline(coef(mod_paired_z)[1], coef(mod_paired_z)[7], col='red')
legend("topleft", legend=c("random", "paired"), fill=c('blue3','red'),
       bty="n", inset=0, cex=.75)
## H
plot(0,0,type="n",xlim=c(6250, 12700), ylim=range(c(paired,random)),
     xaxt="n",xlab="Sampling effort [%]",
     ylab="Nurse effect [z-score]",yaxt="n")
title('H', adj = 0, line = 0)
axis(2,at=c(-20,-10,0,10,20), las=1)
axis(1,at=c(6250, 12500), labels=c(5, 10), las=1)
points(study_df[,7], random, pch = 21, col = 'blue3')
points(study_df[,7]+250, paired, pch = 21, col = 'red')
abline(coef(mod_random_z)[1], coef(mod_random_z)[8], col='blue3')
abline(coef(mod_paired_z)[1], coef(mod_paired_z)[8], col='red')
```

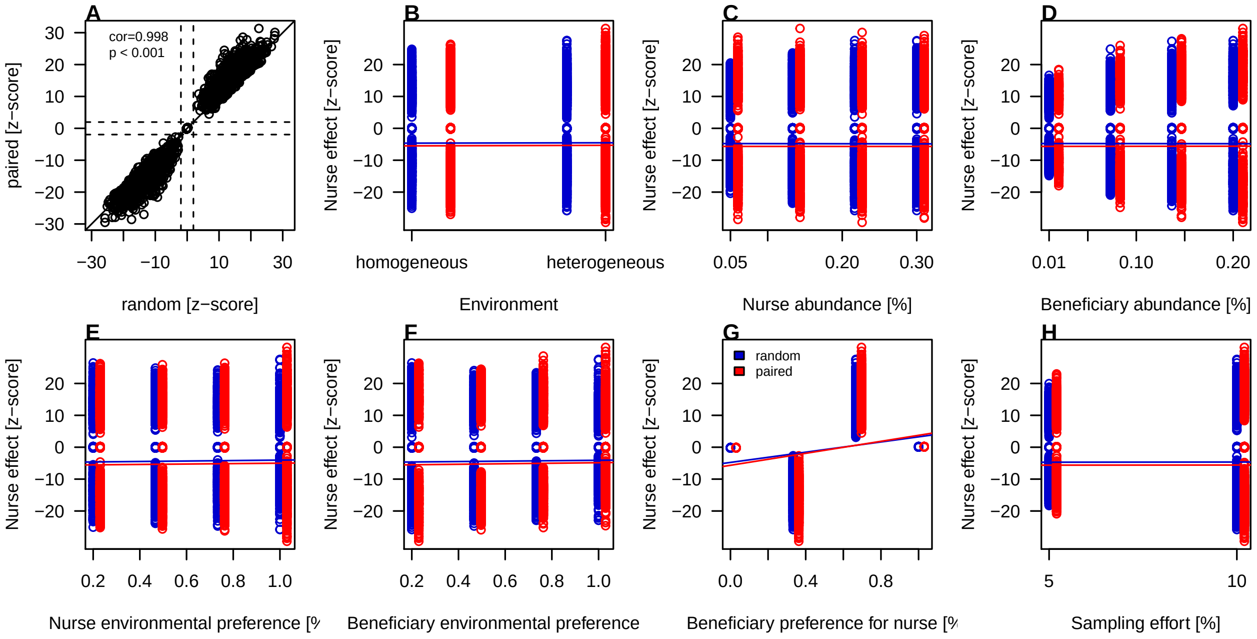

##### Relative differences in parameter estimate $b_1$

```
study_df$delta = (study_df$paired_b-study_df$random_b)/study_df$random_b

study_df$delta = (study_df$paired-study_df$random)

delta = study_df$delta
t.test(study_df$delta)
```

```
##
##  One Sample t-test
##
```

```
## data:  study_df$delta
## t = -1.5848, df = 4095, p-value = 0.1131
## alternative hypothesis: true mean is not equal to 0
## 95 percent confidence interval:
##  -0.12985977  0.01376129
## sample estimates:
##   mean of x
## -0.05804924
```

```
##
mod_delta = lm(delta~ env_h+nur_a+ben_a+nur_p+ben_p+ben_n+sampl,
               data = study_df)
Anova(mod_delta)
```

```
## Anova Table (Type II tests)
##
## Response: delta
##            Sum Sq   Df  F value Pr(>F)
## env_h         2.7    1   0.5110 0.4747
## nur_a         3.3    1   0.6299 0.4274
## ben_a         6.1    1   1.1725 0.2790
## nur_p         1.5    1   0.2888 0.5910
## ben_p         7.3    1   1.3953 0.2376
## ben_n      1084.8    1 207.2516 <2e-16 ***
## sampl         0.4    1   0.0815 0.7753
## Residuals 21396.7 4088
## ---
## Signif. codes:  0 '***' 0.001 '**' 0.01 '*' 0.05 '.' 0.1 ' ' 1
```

```
summary(mod_delta)
```

```
##
## Call:
## lm(formula = delta ~ env_h + nur_a + ben_a + nur_p + ben_p +
##     ben_n + sampl, data = study_df)
##
## Residuals:
##      Min       1Q   Median       3Q      Max
## -13.0369  -0.6613  -0.0583   0.7998  12.7358
##
## Coefficients:
##               Estimate Std. Error t value Pr(>|t|)
## (Intercept) -9.004e-01  1.845e-01  -4.881 1.09e-06 ***
## env_hhetero  5.111e-02  7.149e-02   0.715    0.475
## nur_a        3.045e-01  3.837e-01   0.794    0.427
## ben_a        5.466e-01  5.048e-01   1.083    0.279
```

```
## nur_p       -6.443e-02  1.199e-01  -0.537    0.591
## ben_p        1.416e-01  1.199e-01   1.181    0.238
## ben_n        1.381e+00  9.592e-02  14.396  < 2e-16 ***
## sampl       -3.265e-06  1.144e-05  -0.285    0.775
## ---
## Signif. codes:  0 '***' 0.001 '**' 0.01 '*' 0.05 '.' 0.1 ' ' 1
##
## Residual standard error: 2.288 on 4088 degrees of freedom
## Multiple R-squared:  0.04915,    Adjusted R-squared:  0.04753
## F-statistic: 30.19 on 7 and 4088 DF,  p-value: < 2.2e-16
```

Differences between paired and random are not significantly different from zero. The 95%CI not only includes zero but is also very far from the significance values of -1.96 and 1.96.

With increasing beneficiary preference for the nurse, the paired sampling tend to overestimate the facilitation effect. But, let's remind that this overestimation does not lead to significant differences.

##### Figures

```r
par(mfrow=c(2,4), mar=c(4,4,1.2,1.2))
## A

hist(study_df$delta, las=1, 50, col='green', freq=F,
     xlab= "Paired-random [nurse effect]", main='')
title('A', adj = 0, line = 0)

## B
plot(0,0,type="n",xlim=c(1,2), ylim=range(delta),
     xaxt="n",xlab="Environment",
     ylab="Paired-random [nurse effect]",yaxt="n")
title('B', adj = 0, line = 0)
axis(2,at=c(-10,-5,0,5,10), las=1)
axis(1,at=c(1,2), labels=c('homogeneous','heterogeneous'), las=1)
points(rep(1,casi/2), delta[which(study_df$env_h=='homo')],
       pch = 21, col = 'green')
points(rep(2,casi/2), delta[which(study_df$env_h=='hetero')],
       pch = 21, col = 'green')
abline(coef(mod_delta)[1], coef(mod_delta)[2], col='green')

## C
plot(0,0,type="n",xlim=c(0.05,0.3), ylim=range(delta),
     xaxt="n",xlab="Nurse abundance [%]",
     ylab="Paired-random [nurse effect]",yaxt="n")
title('C', adj = 0, line = 0)
```

```r
axis(2,at=c(-10,-5,0,5,10), las=1)
axis(1,at=c(0.05, 0.1, 0.2, 0.3), las=1)
points(study_df[,2], delta, pch = 21, col = 'green')
abline(coef(mod_delta)[1], coef(mod_delta)[3], col='green')

## D
plot(0,0,type="n",xlim=c(0.01,0.2), ylim=range(delta),
     xaxt="n",xlab="Beneficiary abundance [%]",
     ylab="Paired-random [nurse effect]",yaxt="n")
title('D', adj = 0, line = 0)
axis(2,at=c(-10,-5,0,5,10), las=1)
axis(1,at=c(0.01, 0.05, 0.10, 0.15, 0.20), las=1)
points(study_df[,3], delta, pch = 21, col = 'green')
abline(coef(mod_delta)[1], coef(mod_delta)[4], col='green')

## E
plot(0,0,type="n",xlim=c(0.2,1), ylim=range(delta),
     xaxt="n",xlab="Nurse environmental preference [%]",
     ylab="Paired-random [nurse effect]",yaxt="n")
title('E', adj = 0, line = 0)
axis(2,at=c(-10,-5,0,5,10), las=1)
axis(1,at=c(0.2, 0.4, 0.6, 0.8, 1.0), las=1)
points(study_df[,4], delta, pch = 21, col = 'green')
abline(coef(mod_delta)[1], coef(mod_delta)[5], col='green')

## F
plot(0,0,type="n",xlim=c(0.2,1), ylim=range(delta),
     xaxt="n",xlab="Beneficiary environmental preference [%]",
     ylab="Paired-random [nurse effect]",yaxt="n")
title('F', adj = 0, line = 0)
axis(2,at=c(-10,-5,0,5,10), las=1)
axis(1,at=c(0.2, 0.4, 0.6, 0.8, 1.0), las=1)
points(study_df[,5], delta, pch = 21, col = 'green')
abline(coef(mod_delta)[1], coef(mod_delta)[6], col='green')

## G
plot(0,0,type="n",xlim=c(0,1), ylim=range(delta),
     xaxt="n",xlab="Beneficiary preference for nurse [%]",
     ylab="Paired-random [nurse effect]",yaxt="n")
title('G', adj = 0, line = 0)
axis(2,at=c(-10,-5,0,5,10), las=1)
axis(1,at=c(0, 0.2, 0.4, 0.6, 0.8, 1.0), las=1)
points(study_df[,6], delta, pch = 21, col = 'green')
abline(coef(mod_delta)[1], coef(mod_delta)[7], col='green')
```

```r
## H
plot(0,0,type="n",xlim=c(6250, 12500), ylim=range(delta),
     xaxt="n",xlab="Sampling effort [%]",
     ylab="Paired-random [nurse effect]",yaxt="n")
title('H', adj = 0, line = 0)
axis(2,at=c(-10,-5,0,5,10), las=1)
axis(1,at=c(6250, 12500), labels=c(5, 10), las=1)
points(study_df[,7], delta, pch = 21, col = 'green')
abline(coef(mod_delta)[1], coef(mod_delta)[8], col='green')
```

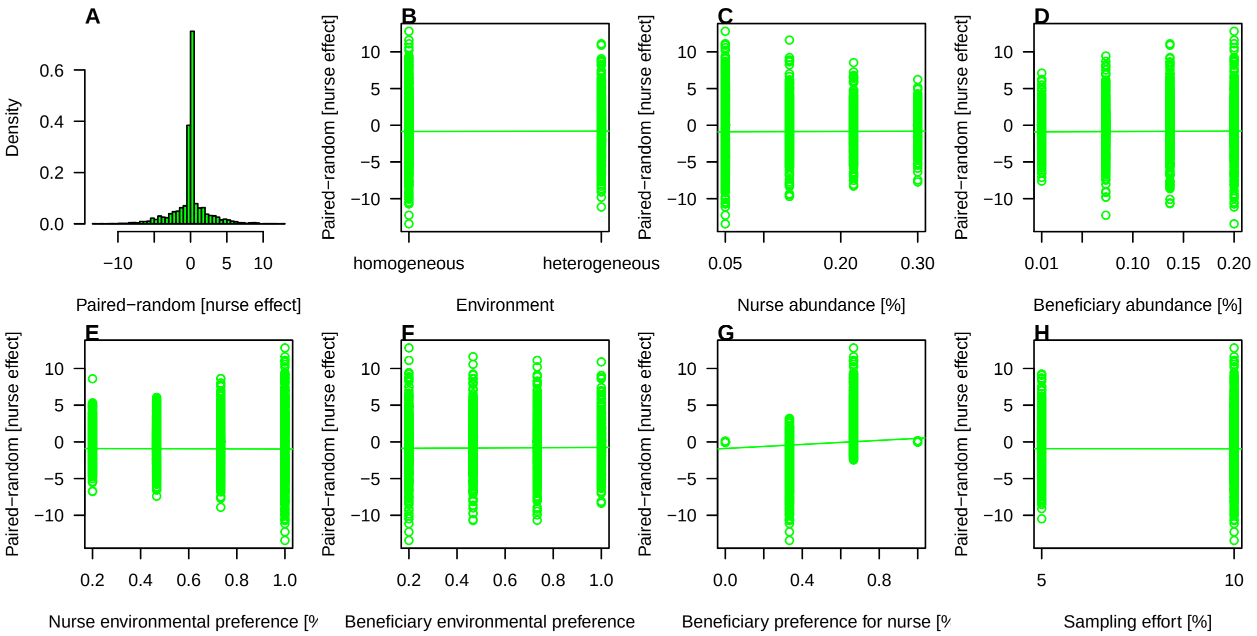

#### Spatial grid

We then simulated communities with a different spatial structure of environmental variation. We made use of the codispersion model they suggested using the R code from Buckley et al. 2016 (Ecology 97, 32–39).

##### Codispersion function

```r
# code from Buckley et al. 2016 Ecology, 97(1): 32-39 [see SI, Appendix]
## Function to simulate co-occurrence patterns on grids

#    grid.points=20       # scale: grain of grid
#    sp1.pattern="CSR"    # The desired pattern for species 1
#    sp2.pattern="CSR"    # The desired pattern for species 2
#    xmin=0               # Dimensions of the plot area
#    xmax=300
```

```
#   ymin=0
#   ymax=300
#   Print=TRUE          # Whether you want plots of the spatial patterns or not

copp.fn <- function(grid.points = grid.points,
                    sp1.pattern = c("CSR","decreasing.x","increasing.x",
                                    "decreasing.y","decreasing.xy","increasing.xy",
                                    "bivariate.normal"),
                    sp2.pattern = c("CSR","decreasing.x","increasing.x",
                                    "increasing.y","decreasing.xy","increasing.xy",
                                    "bivariate.normal","inc.x.dec.y","dec.x.inc.y"),
                    xmin=xmin,xmax=xmax,ymin=ymin,ymax=ymax,
                    Print=c("TRUE","FALSE")){

  # 1. Create an empty list to add output geodata objects
  copp.sim <- vector("list",2)

  # 2. Set up underlying grid coordinates
  X <- seq(from=xmin,to=xmax-grid.points,by=grid.points)
  Y <- seq(from=ymin,to=ymax-grid.points,by=grid.points)
  gridxy <- expand.grid(x=X,y=Y)

  # 3a. Create a set of quadrat abundance values for sp1
  # based on the 'sp1.pattern' argument

  if(sp1.pattern=="CSR"){Z <- rnorm(length(gridxy$x),mean=30,sd=10) }

  if(sp1.pattern=="decreasing.x"){Z <- 1+(rev(2*gridxy$x+5))/10 }

  if(sp1.pattern=="increasing.x"){Z <- 1+(2*gridxy$x+5)/10 }

  if(sp1.pattern=="decreasing.y"){Z <- 1+(rev(2*gridxy$y+5))/10 }

  if(sp1.pattern=="decreasing.xy"){
    Z <- 1+rev(((gridxy$x+1)^2+(gridxy$y+1)^2)/3000) # (x-u)^2+(y-v)^2
  }

  if(sp1.pattern=="increasing.xy"){
    Z <- 1+((gridxy$x+2)^2+(gridxy$y+1)^2)/3000 # (x-u)^2+(y-v)^2
  }

  if(sp1.pattern=="bivariate.normal"){
    Z <- 300*bivariate(((gridxy$x-min(gridxy$x))/
                          (max(gridxy$x)-min(gridxy$x))*4)-2,((gridxy$y-min(gridxy$y))
```

```
                                                                  (max(gridxy$y)-min(g
} #  bivariate normal

if(sp1.pattern=="inc.x.dec.y"){
  Z <- 1+((gridxy$x+2)^2+(rev(gridxy$y+1))^2)/3000 # (x-u)^2+(y-v)^2
}

if(sp1.pattern=="dec.x.inc.y"){
  Z <- 1+((rev(gridxy$x+2))^2+(gridxy$y+1)^2)/3000 # (x-u)^2+(y-v)^2
}

# 3b. Add data to the output list as a geodata object
copp.sp1.df <- data.frame(x=gridxy$x,y=gridxy$y,Z=jitter(Z,mean(Z)/5))
copp.sim[[1]] <- as.geodata(copp.sp1.df,coords.col=1:2,data.col=3)

# 4a. Create a set of quadrat abundance values for sp2
# based on the 'sp1.pattern' argument

if(sp2.pattern=="CSR"){Z <- rnorm(length(gridxy$x),mean=30,sd=10) }

if(sp2.pattern=="decreasing.x"){Z <- 1+(rev(2*gridxy$x+5))/10 }

if(sp2.pattern=="decreasing.x"){Z <- 1+(rev(2*gridxy$x+5))/10 }

if(sp2.pattern=="increasing.x"){Z <- 1+(2*gridxy$x+5)/10 }

if(sp2.pattern=="increasing.y"){Z <- 1+(2*gridxy$y+5)/10 }

if(sp2.pattern=="decreasing.xy"){
  Z <- 1+rev(((gridxy$x+1)^2+(gridxy$y+1)^2)/3000) # (x-u)^2+(y-v)^2
}

if(sp2.pattern=="increasing.xy"){
  Z <- 1+((gridxy$x+2)^2+(gridxy$y+1)^2)/3000 # (x-u)^2+(y-v)^2
}

if(sp2.pattern=="bivariate.normal"){
  Z <- 300*bivariate(((gridxy$x-min(gridxy$x))/
                          (max(gridxy$x)-min(gridxy$x))*4)-2,
                      ((gridxy$y-min(gridxy$y))/
                                                                  (max(gridxy$y)-min(g
} #  bivariate normal

if(sp2.pattern=="inc.x.dec.y"){
```

```
    Z <- 1+((gridxy$x+2)^2+(rev(gridxy$y+1))^2)/3000 # (x-u)^2+(y-v)^2
  }

  if(sp2.pattern=="dec.x.inc.y"){
    Z <- 1+((rev(gridxy$x+2))^2+(gridxy$y+1)^2)/3000 # (x-u)^2+(y-v)^2
  }

  # 4b. Add data to the output list as a geodata object
  copp.sp2.df <- data.frame(x=gridxy$x,y=gridxy$y,Z=jitter(Z,mean(Z)/10))
  copp.sim[[2]] <- as.geodata(copp.sp2.df,coords.col=1:2,data.col=3)

  # 5. Print map of points if desired
  if(Print=="TRUE"){
    print(qplot(x, y, data=copp.sp1.df, size=Z,
                main=paste("sp1.pattern = ",sp1.pattern))+ theme_bw())
    print(qplot(x, y, data=copp.sp2.df, size=Z,
                main=paste("sp2.pattern = ",sp2.pattern))+ theme_bw())
    } # end Print loop

  # 6. Output the list of geodata objects
  return(copp.sim)
  } # end function
```

##### Simulation

We generated a bivariate spatial pattern with an abundance gradient decreasing from bottom to top. This simulation was replicated 50 times.

```
## Run the simulation
set.seed(20211011)

## list with models
mods99 = rep(list(NA),99)
## run models
for(i in 1:99){
    print(i)
mods99[[i]] = copp.fn(grid.points=5,
                      sp1.pattern="decreasing.y",
                      sp2.pattern="decreasing.y",
                      xmin=0, xmax=500, ymin=0, ymax=500,
                      Print=FALSE)
}

## dataframe with parms
parmsdf = data.frame(n=rep(1:50,2),
```

```r
                     parm = rep(NA,50*2),
                     tval = rep(NA,50*2),
                     method = gl(2,50,2*50)
                     )

## parameters random sampling
mod_rs <- foreach(i=1:50, .inorder = TRUE) %dopar% {
mod1=mods99[[i]]
## random sampling
rs1=sample(1:10000, 500)
rs1=sort(rs1)
rs2=sample(1:10000, 500)
rs2=sort(rs2)
dataf.rs=data.frame(rs_sp1= mod1[[1]]$data[rs1],
                    rs_sp2= mod1[[2]]$data[rs2],
                    dy= sqrt(mod1[[1]][[1]][,2][rs1]*
                                mod1[[2]][[1]][,2][rs2])+
                                rnorm(500,0,0.1)
                    )
gls(rs_sp2 ~ rs_sp1, data = dataf.rs, correlation=corLin(form=~dy))
}

for(i in 1:50){
    parmsdf[i,2]=summary(mod_rs[[i]])[[18]][2,1]
    parmsdf[i,3]=summary(mod_rs[[i]])[[18]][2,3]
}

## paired sampling
mod_ps <- foreach(i=1:50, .inorder = TRUE) %dopar% {
mod1=mods99[[i]]
pairsamp = sample(1:10000, 500)
dataf.ps=data.frame(rs1=rs1,
                    dy= mod1[[1]][[1]][,2][pairsamp]+rnorm(500,0,0.1),
                    ps_sp1= mod1[[1]]$data[pairsamp],
                    ps_sp2= mod1[[2]]$data[pairsamp])
gls(ps_sp2 ~ ps_sp1, data = dataf.ps, correlation=corLin(form=~ dy))
}

for(i in 1:50){
    parmsdf[i+50,2]=summary(mod_ps[[i]])[[18]][2,1]
    parmsdf[i+50,3]=summary(mod_ps[[i]])[[18]][2,3]
}
```

#### Statistical analysis of simulated plant communities

```
mod2.parm = lme(parm ~ method, data=parmsdf, random = ~1|n)
summary(mod2.parm)

mod2.tval = lme(tval ~ method, data=parmsdf, random = ~1|n)
summary(mod2.tval)

### false negatives, tipe-2 error rate
## among random
length(which(parmsdf[1:50,3]<1.96))/50*100
parmsdf[which(parmsdf[1:50,3]<1.96),3]

## among paired
length(which(parmsdf[51:1000,3]<1.96))/50*100

### percentage
0.1013711/0.8970443*100
193.2144/296.5086*100

### variance
sd(parmsdf[1:50,2])
sd(parmsdf[1:50,3])
sd(parmsdf[51:100,2])
sd(parmsdf[51:100,3])
```

#### Figure

```
boxplot(tval ~ method, data=parmsdf, las = 1,
        col = c('lightblue','green'),
        xlab=c('Random', 'Paired'), ylab = 't-value')
```

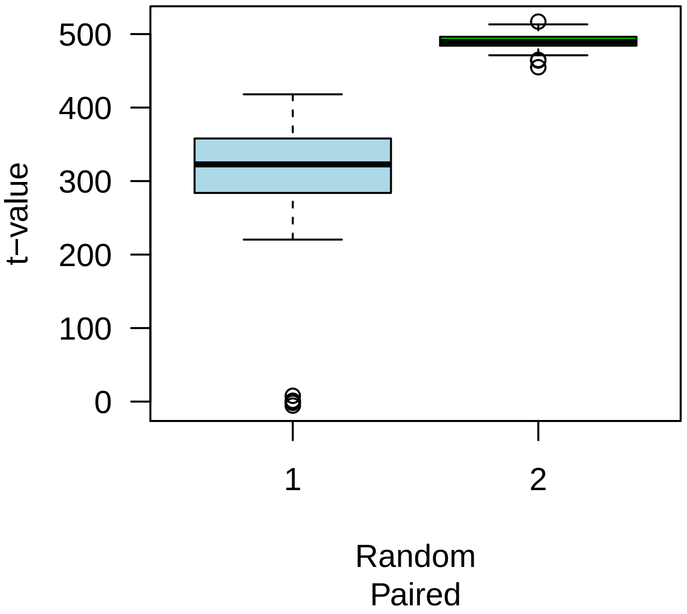
t−value
500
400
300
200
100
0
1
2
Random
Paired
