## Supplementary Information Talbe for "Assessing the accuracy of paired and random sampling for quantifying plant–plant interactions in natural communities"

**Table S1.** Type-II-ANOVA of the linear model testing the effects of habitat quality, nurse abundance, beneficiary abundance, nurse habitat preference, beneficiary habitat preference, beneficiary affinity to nurse, sampling effort as well as sampling method alone and in interactions with all these previous factors on facilitation effects (*z-*score).

| Factor | DF | Sum Sq | F-value | P-value |  |
| --- | --- | --- | --- | --- | --- |
| Method | 1 | 7 | 0.07 | 0.793 |  |
| Habitat quality | 1 | 43 | 0.43 | 0.513 |  |
| Nurse abundance | 1 | 2 | 0.02 | 0.876 |  |
| Beneficiary abundance | 1 | 0 | 0.00 | 0.998 |  |
| Nurse habitat preference | 1 | 351 | 3.51 | 0.061 | . |
| Beneficiary habitat preference | 1 | 399 | 3.99 | 0.0456 | * |
| Beneficiary affinity to nurse | 1 | 86359 | 865.7 | <0.001 | *** |
| Sampling effort | 1 | 6 | 0.06 | 0.813 |  |
| Method × Habitat quality | 1 | 1 | 0.01 | 0.908 |  |
| Method × Nurse abundance | 1 | 2 | 0.02 | 0.898 |  |
| Method × Beneficiary abundance | 1 | 3 | 0.03 | 0.861 |  |
| Method × Nurse habitat preference | 1 | 1 | 0.01 | 0.931 |  |
| Method × Beneficiary habitat preference | 1 | 4 | 0.04 | 0.848 |  |
| Method × Beneficiary affinity to nurse | 1 | 542 | 5.43 | 0.020 | * |
| Method × Sampling effort | 1 | 0 | 0.00 | 0.963 |  |
| *Residuals* | 8176 | 815608 |  |  |  |

**Table S2.** Summary of the linear model testing the effects of habitat quality, nurse abundance, beneficiary abundance, nurse habitat preference, beneficiary habitat preference, beneficiary affinity to nurse, sampling effort as well as sampling method alone and in interactions with all these previous factors on facilitation effects (*z-*score). Multiple R-squared = 0.09711, Adjusted R-squared = 0.09545. F-statistic: 58.62 on 15 and 8176 DF, P-value <0.001.

| Coefficient | Beta | SE | t-value | P-value |  |
| --- | --- | --- | --- | --- | --- |
| Intercept | -4.77 | 0.81 | -5.92 | <0.001 |  |
| Method | -0.90 | 1.14 | -0.79 | 0.429 |  |
| Habitat quality | 0.12 | 0.31 | 0.38 | 0.704 |  |
| Nurse abundance | -0.34 | 1.68 | -0.20 | 0.841 |  |
| Beneficiary abundance | -0.27 | 2.20 | -0.12 | 0.903 |  |
| Nurse habitat preference | 0.73 | 0.52 | 1.39 | 0.165 |  |
| Beneficiary habitat preference | 0.67 | 0.52 | 1.28 | 0.201 |  |
| Beneficiary affinity to nurse | 8.02 | 0.42 | 19.16 | <0.001 | *** |
| Sampling effort | <0.01 | <0.01 | 0.20 | 0.841 |  |
| Method × Habitat quality | 0.05 | 0.44 | 0.12 | 0.908 |  |
| Method × Nurse abundance | 0.30 | 2.37 | 0.13 | 0.898 |  |
| Method × Beneficiary abundance | 0.55 | 3.12 | 0.18 | 0.861 |  |
| Method × Nurse habitat preference | 0.06 | 0.74 | -0.09 | 0.931 |  |
| Method × Beneficiary habitat preference | 0.14 | 0.74 | 0.19 | 0.848 |  |
| Method × Beneficiary affinity to nurse | 1.38 | 0.59 | 2.33 | 0.020 | * |
| Method × Sampling effort | <0.01 | <0.01 | -0.05 | 0.963 |  |
| *Residuals* |  | 9.99 |  |  |  |

**Table S3.** Type-I-ANOVA of the linear mixed effects model testing the effects of the stress level, sampling method, neighbour presence and beneficiary species identity on the abundance of the two beneficiaries at the Spanish site.

| Factor | nomDF | den.DF | F-value | P-value |  |
| --- | --- | --- | --- | --- | --- |
| (Intercept) | 1 | 16.3 | 327.70 | <0.001 | *** |
| Stress | 2 | 37.0 | 20.93 | <0.001 | *** |
| Method | 1 | 55.9 | 1.93 | 0.17 |  |
| Neighbour | 1 | 94.0 | 0.50 | 0.48 |  |
| Beneficiary | 1 | 550.1 | 134.20 | <0.001 | *** |
| Stress × Method | 2 | 56.1 | 1.25 | 0.30 |  |
| Stress × Neighbour | 2 | 92.9 | 2.37 | 0.099 | . |
| Method × Neighbour | 1 | 94.7 | 0.60 | 0.44 |  |
| Stress × Beneficiary | 2 | 550.1 | 31.05 | <0.001 | *** |
| Method × Beneficiary | 1 | 550.1 | 3.86 | 0.050 | . |
| Neighbour × Beneficiary | 1 | 550.1 | 22.52 | <0.001 | *** |
| Stress × Method × Neighbour | 2 | 96.3 | 1.39 | 0.26 |  |
| Stress × Method × Beneficiary | 2 | 550.1 | 1.94 | 0.14 |  |
| Stress × Neighbour × Beneficiary | 2 | 550.1 | 3.47 | 0.032 | * |
| Method × Neighbour × Beneficiary | 1 | 550.1 | 0.00 | 0.96 |  |
| Stress × Method × Neighbour × Ben. | 2 | 550.1 | 0.25 | 0.78 |  |
| Variance components: |  |  |  |  |  |
| Random factor | Var. | SE | z.ratio |  |  |
| Replicate | -41524 | 112766 | -0.37 |  |  |
| Stress × Replicate | -156470 | 287643 | -0.54 |  |  |
| Stress × Method × Replicate | 1013273 | 423127 | 2.39 |  |  |
| Stress × Method × Neigh. × Rep. | -532255 | 303331 | -1.75 |  |  |
| Residuals | 9017491 | 543747 | 16.58 |  |  |
| Dispersion: 3002.91 |  |  |  |  |  |

**Table S4.** Type-I-ANOVA of the linear mixed effects model testing the effects of the sampling method, neighbour presence, canopy position and beneficiary species identity on the abundance of the two beneficiary species at the Italian site.

| Factor | nomDF | denomDF | F-value | P-value |  |
| --- | --- | --- | --- | --- | --- |
| (Intercept) | 1 | 14.5 | 382.20 | <0.001 | *** |
| Method | 1 | 15.1 | 0.59 | 0.45 |  |
| Neighbour | 1 | 29.7 | 54.67 | <0.001 | *** |
| Canopy position | 2 | 113.8 | 41.96 | <0.001 | *** |
| Beneficiary | 1 | 526.7 | 17.11 | <0.001 | *** |
| Method × Neighbour | 1 | 29.6 | 0.95 | 0.34 |  |
| Method × Canopy position | 2 | 114.0 | 0.65 | 0.52 |  |
| Neighbour × Canopy position | 2 | 114.2 | 34.96 | <0.001 | *** |
| Method × Beneficiary | 1 | 526.7 | 10.70 | 0.001 | ** |
| Neighbour × Beneficiary | 1 | 526.7 | 193.50 | <0.001 | *** |
| Canopy position × Beneficiary | 2 | 526.7 | 11.64 | <0.001 | *** |
| Method × Neighbour × Canopy position | 2 | 114.5 | 4.47 | 0.01 | * |
| Method × Neighbour × Beneficiary | 1 | 526.7 | 2.09 | 0.15 |  |
| Method × Canopy position × Beneficiary | 2 | 526.7 | 0.18 | 0.83 |  |
| Neighbour × Canopy position × Ben. | 2 | 526.7 | 10.27 | <0.001 | *** |
| Method × Neighbour × C. position × Ben. | 2 | 526.7 | 0.53 | 0.59 |  |
| Variance components: |  |  |  |  |  |
| Random factor | Variance | SE | z.ratio |  |  |
| Replicate | -0.27 | 0.41 | -0.67 |  |  |
| Method × Replicate | -0.09 | 0.83 | -0.11 |  |  |
| Neighbour × Method × Replicate | 2.81 | 1.00 | 2.82 |  |  |
| C. position × Neighbour × Method × Replicate | -3.21 | 0.50 | -6.38 |  |  |
| Residuals | 23.23 | 1.43 | 16.23 |  |  |
| Dispersion: 4.82 |  |  |  |  |  |

**Table S5.** Type-I-ANOVA of the linear mixed effects model testing the effects of the sampling method and neighbour presence on the species richness of the community (upper table) and frequency of occurrence of *Salix nummularia* (lower table) at the Japanese site.

| Species richness | (square-root transformed) | | |  |  |  |
| --- | --- | --- | --- | --- | --- | --- |
| Factor | df | SS | MS | F-value | P-value |  |
| Method | 1 | 0.02 | 0.02 | 0.03 | 0.858 |  |
| Neighbour | 1 | 21.19 | 21.19 | 36.52 | <0.001 | *** |
| Method X Neighbour | 1 | 0.31 | 0.31 | 0.53 | 0.468 |  |
| Residuals | 244 | 141.57 | 0.58 |  |  |  |

| *Salix nummularia* | |  |  |  |  |
| --- | --- | --- | --- | --- | --- |
| Factor | df | deviance | residual df | residual deviance | P-value |
| Method | 1 | 8.67 | 246 | 243.36 | 0.003 |
| Neighbour | 1 | 48.72 | 245 | 194.64 | <0.001 |
| Method × Neighbour | 1 | 0.35 | 244 | 194.29 | 0.556 |
